# Generating realistic neurophysiological time series with denoising diffusion probabilistic models

**DOI:** 10.1101/2023.08.23.554148

**Authors:** Julius Vetter, Jakob H. Macke, Richard Gao

## Abstract

In recent years, deep generative models have had a profound impact in engineering and sciences, revolutionizing domains such as image and audio generation, as well as advancing our ability to model scientific data. In particular, Denoising Diffusion Probabilistic Models (DDPMs) have been shown to accurately model time series as complex high-dimensional probability distributions. Experimental and clinical neuroscience also stand to benefit from this progress, since accurate modeling of neurophysiological time series, such as electroencephalography (EEG), electrocorticography (ECoG), and local field potential (LFP) recordings, and their synthetic generation can enable or improve a variety of neuroscientific applications. Here, we present a method for modeling multi-channel and densely sampled neurophysiological recordings using DDPMs, which can be flexibly applied to different recording modalities and experimental configurations. First, we show that DDPMs can generate realistic synthetic data for a variety of datasets including different recording techniques (LFP, ECoG, EEG) and species (rat, macaque, human). DDPM-generated time series accurately capture single- and multi-channel statistics such as frequency spectra and phase-amplitude coupling, as well as fine-grained and dataset-specific features such as sharp wave-ripples. In addition, synthetic time series can be generated based on additional information like experimental conditions or brain states. We demonstrate the utility and flexibility of DDPMs in several neuroscience-specific analyses, such as brain-state classification and imputation of missing channels to improve neural decoding. In summary, DDPMs can serve as accurate generative models of neurophysiological recordings, and have a broad utility in the probabilistic generation of synthetic time series for neuroscientific applications.

## 1 Introduction

Statistical models that can accurately reconstruct complex datasets have long been useful in many areas of science. These so-called generative models allow scientists to interpret and analyze statistical dependencies between data features, as well as between data and mechanistic latent variables. Such models further grant the ability to generate realistic synthetic data, which is valuable in its own right, especially when it is possible to incorporate (or “condition on”) additional information or observations into the model. Examples of applications include imputing missing data given observed data, forecasting the future given past data, as well as creating synthetic datasets as input to a simulator and to replace or augment scarce training data for subsequent machine learning models, all of which rely on a (conditional) generative model that can produce realistic synthetic data.

The generation of synthetic neurophysiological recordings has received a great deal of attention over the years, predominantly with a focus on generating spike trains, i.e., binary sequences of action potential times: Krumin and Shoham (2009), Macke et al. (2009), Gutnisky and Josić (2010) use thresholding of Gaussian processes and similar techniques to generate spike trains with a pre-specified correlation structure. More generally, time series surrogate methods (Schreiber and Schmitz, 2000, Venema et al., 2006) can be used to generate synthetic neurophysiological recordings that mimic the real data in a well-defined set of features, but lose complex, nonlinear features by design. Other works have employed deep neural networks as non-linear encoding models of brain response, e.g., predicting EEG (Gifford et al., 2022) response from viewed images. However, they are deterministic by design and cannot be used to probabilistically simulate neural time series, nor can they be used without conditioning information such as visual stimuli.

Recently, generative models that leverage deep neural networks for probabilistic modeling have made a significant impact across various domains, transforming the way we approach tasks such as image and audio generation (Kong et al., 2021, Dhariwal and Nichol, 2021, Rombach et al., 2022), time series imputation (Fortuin et al., 2020), as well as in scientific applications ranging from molecular design (Sanchez-Lengeling and Aspuru-Guzik, 2018) to black hole imaging (Sun and Bouman, 2021). Neuroscience research, in particular, has benefited from various types of deep generative models. For example, variational autoencoders (VAEs) have been applied to infer low-dimensional representations of single-trial neural population dynamics (Pandarinath et al., 2018), while generative adversarial networks (GANs) have been used for the task of spike-train generation (Molano-Mazon et al., 2018, Ramesh et al., 2019), as well as to decode images from single neuron and fMRI data (Ponce et al., 2019, Lin et al., 2022). The recently proposed denoising diffusion probabilistic models (DDPMs) have also been applied to improve neural decoding performance, in particular leveraging latent diffusion models (Rombach et al., 2022) to predict viewed images from fMRI data (Takagi and Nishimoto, 2022, Chen et al., 2023).

Despite their utility for encoding and decoding spiking and imaging data, deep generative models have not yet found empirical success in realistic reproduction of a type of data that is ubiquitous in neuroscience: multivariate and densely sampled electrophysiological time series. Such recordings include non-invasive scalp electroencephalography (EEG), as well as intracranial electrocorticography (ECoG) and local field potential (LFP), which are routinely recorded during scientific research and clinical brain monitoring.

A general-purpose model that can conditionally synthesize such neurophysiological data would be valuable in many scenarios. For example, brain recordings often contain missing values due to sensor noise or misplacement (Silva et al., 2012), which limits their use in neuroscience-specific applications and can lead to them being discarded altogether. Such practical problems are exacerbated by the fact that individual neurophysiological datasets are often limited in availability due to restrictions in clinical contexts. Deep generative models for time series can partially alleviate these issues by providing synthetic data, for example, to facilitate the training of a brain-computer interface (BCI) or as input data for scientific simulations with reduced privacy concerns (Walonoski et al., 2018, Yoon et al., 2019, Abbasi et al., 2020). However, while deep generative models and diffusion models in particular have been used for modeling complex high-dimensional time series in a number of settings (Lin et al., 2023), including probabilistic imputation (Tashiro et al., 2021, Alcaraz and Strodthoff, 2022) and forecasting (Rasul et al., 2021, Biloš et al., 2022), their general utility and performance in modeling neurophysiological recordings have thus far been unexplored.

In this work, we demonstrate that denoising diffusion probabilistic models (DDPMs) (Ho et al., 2020)—or diffusion models for short—can accurately generate multivariate and densely sampled continuous neurophysiological recordings, and further showcase their utility in a variety of neuroscience-specific applications. In particular, we use deep neural networks with structured convolutions (Li et al., 2023) and Ornstein-Uhlenbeck (OU) processes as diffusion processes (Biloš et al., 2022) to improve the modeling of such time series with DDPMs, which allows trained models to produce synthetic recordings with realistic power spectra and even complex features such as sharp wave-ripples and cross-frequency coupling. By conditioning on observed values and other behavioral or experimental variables, synthetic recordings automatically capture cross-area interaction between many channels, which is used to impute missing channels and improve performance in a human BCI application at test time. Finally, DDPMs provide additional benefits accessible to generative models, enabling behavioral state classification and outlier detection without additional network training. Code for all our experiments is available at https://github.com/mackelab/neural_timeseries_diffusion.

## 2 Results

### 2.1 Denoising Diffusion Probabilistic Models for neurophysiological time series

We use DDPMs (Ho et al., 2020) to learn generative models of electrophysiological recordings. DDPMs are defined by a forward (noising) process and a reverse (denoising) process. The noising process successively adds random noise from a pre-specified distribution (usually independent Gaussian) to the real data through a series of diffusion steps. The reverse process uses a deep neural network as a “denoiser”, which is trained to reverse the forward process at each diffusion step (Fig. 1A). To generate synthetic data, samples of random Gaussian noise are drawn and gradually denoised by the trained network (Fig. 1A box, top to bottom) over the same number of diffusion steps, resulting in synthetic data that follows the statistical distribution of the real data (i.e., unconditional generation or simulation, Fig. 1B). The training and inference procedure can also incorporate additional information through conditioning. For example, when real data from other channels is given alongside the initial noise sample for the generated channel, the denoising process is guided to generate synthetic data that is likely given the other channels, which can be used to perform imputation of missing channels or interpolation (Fig. 1C). Similarly, when behavioral or experimental state variables are included during training, they can be used to guide the denoising process during time series generation (Fig. 1D). In addition, such a conditionally trained model can also act as a classifier or outlier detector, since the model can be queried for the likelihood of observing a data sample given either conditions.

**Figure 1:**
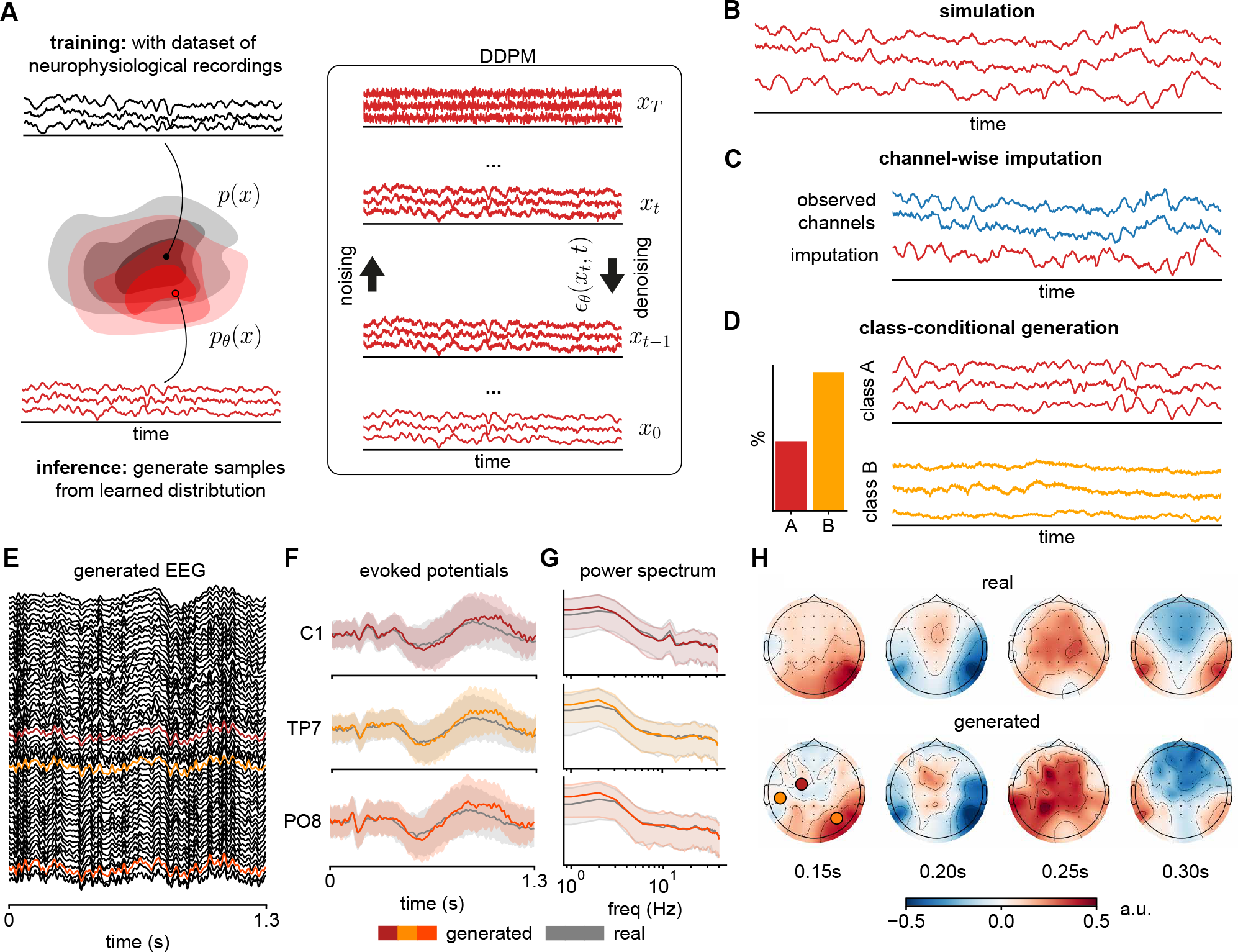
Overview of diffusion models for neurophysiological recordings and subsequent applications, and an example of DDPM-generated EEG signal. **(A)** A denoising diffusion probabilistic model (DDPM) *p*_*θ*_(*x*) is trained on a dataset of neurophysiological recordings. It attempts to generate samples from the data distribution *p*(*x*), underlying the training data, by successively denoising samples from a pre-specified Gaussian distribution, using a neural network ***ϵ***_*θ*_(*x*_*t*_, *t*) as the denoiser. In the context of neurophysiological recordings, DDPMs can be used for various different tasks. Examples include **(B)** simulation of neurophysiological recordings, **(C)** imputation of missing values in these recordings, and **(D)** class-conditional generation of recordings from different experimental conditions or brain states. Since DDPMs allow the computation of likelihoods, the class-conditional model can also be used to perform tasks like classification or outlier detection. **(E)** An example DDPM-generated trial of 56-channel EEG. **(F)** Trial-average of three channels show close overlap between real (gray) and generated (colored) evoked potentials (mean and standard deviation across trials), **(G)** power spectra (median and 10% / 90% percentiles), and **(H)** spatio-temporal relationships reflected in scalp topography.

We follow the standard DDPM training and inference procedure (Ho et al., 2020), while incorporating two recent innovations in time series modeling: First, since neurophysiological time series are densely sampled (for example, sampling rate of 500 Hz or more), learning long-range dependencies over hundreds of time points in just one second of data is a non-trivial challenge. We address this problem by using structured convolutions in the denoising network, which use a collection of multi-scale convolution kernels that have shown performance and efficiency improvements on long-range time series tasks (Li et al., 2023). Second, neurophysiological recordings of all modalities (EEG, ECoG, LFP) follow a 1*/f* -like power law in the frequency domain under a range of behavioral and brain states (Pritchard, 1992, He et al., 2010, Gao et al., 2017, Donoghue et al., 2020), in contrast to the flat spectrum of the Gaussian white noise commonly used in DDPMs. Therefore, we sometimes use the Ornstein-Uhlenbeck (OU) process (i.e., colored noise) in the forward noising process (Biloš et al., 2022) to incorporate this prior knowledge, and empirically demonstrate improvements over the standard white noise in a number of experiments. Full details on DDPMs, as well as our denoising network architecture and parameterization of the OU process can be found in Methods (Section 4).

As a first demonstration, we train a diffusion model on a single participant’s EEG recorded during a trial-structured task (BCI Challenge @ NER 2015, Margaux et al. (2012)). The trained DDPM is able to simulate (i.e., unconditionally generate) realistic single-trial EEG data (Fig. 1E) and capture features in the real trial-averaged time series and power spectra (Fig. 1F, G), in particular the evolution of the evoked potentials, and a 12 Hz oscillation in the C1-channel. In addition, spatio-temporal relationships across the entire scalp are preserved, as shown in the topographic plots over time (Fig. 1H). This introductory demonstration illustrates how DDPMs can realistically generate high-dimensional neurophysiological time series, and in the following sections we further evaluate and exploit this capability on a variety of datasets and applications.

### 2.2 Overview of DDPM applications: experiments and datasets

Beyond the first example, we apply our model to three different datasets to test whether diffusion models can generate realistic synthetic brain recordings in different settings, and to demonstrate their utility in a variety of tasks relevant to neuroscience. On all three datasets, DDPMs accurately capture the statistics of real recordings—in particular, the distribution of channel-wise power spectral densities (PSDs), as well as the multivariate cross-spectrum—across species, recording modalities, and brain areas. Critically, the trained models do not simply overfit to or memorize samples from the training set, which we confirm by measuring the pairwise distance between generated and training data (see Appendix). We show that DDPMs are a general-purpose generative model that accurately captures idiosyncratic features in each of the three datasets, and can be applied in a variety of tasks with minor modifications to network architecture or training procedure: The first dataset consists of 3-channel local field potential (LFP) recordings from rat prefrontal cortex, thalamus, and hippocampus (Varela and Wilson, 2020). These recordings contain sharp wave-ripples that exhibit complex cross-frequency and cross-channel dependencies, which are correctly captured by our model in both simulation and imputation settings. The second dataset consists of human electrocorticography (ECoG) recorded during naturalistic behavior from 12 participants undergoing epilepsy monitoring (AJILE12, Peterson et al. (2022)). In addition to accurately generating high channel-count ECoG data, we show that DDPMs can sensibly impute corrupted or missing channels in an artificially induced missing-data setting (similar to Talukder et al. (2022)), which substantially improves performance in a neural decoding task compared to a baseline (zero-)imputation. The third dataset consists of whole-brain ECoG recordings from a macaque monkey under two different brain states (awake and anesthetized). DDPMs can conditionally generate recordings given one of these two states, as well as evaluate class-specific likelihoods, which in turn can be used to perform classification and outlier detection. The rest of this section presents detailed results from each of the three experiments.

### 2.3 DDPM captures cross-frequency and cross-region dependencies of sharp wave-ripples

In the first experiment, we apply our DDPM to model LFP recorded from freely behaving rats (sampled at 600 Hz) in three locations: medial prefrontal cortex (mPFC), hippocampal CA1 region, and the thalamic nucleus reuniens (RE) (Varela and Wilson, 2019). These recordings contain neurophysiological features of interest, such as slow oscillations, spindles, and sharp wave-ripples (SWR), which are region-specific and occur with a specific phase relationship relative to one another (Varela and Wilson, 2020). Thus, we aim to generate synthetic data and assess whether our model is able to correctly capture the single-channel voltage distribution and power spectrum, but also complex features such as phase-amplitude coupling between slow oscillations and ripples across channels.

After training the model on 3480 2-second time series, we generate the same number of synthetic samples for comparison. Visually, generated traces look indistinguishable from real data, and contain clearly visible SWRs (example in Fig. 2B). Across all generated samples, the distribution of voltage values and power spectral density between real and synthetic data match closely in all three regions (Fig. 2C), and components such as 60 Hz line-noise are also reproduced correctly.

**Figure 2:**
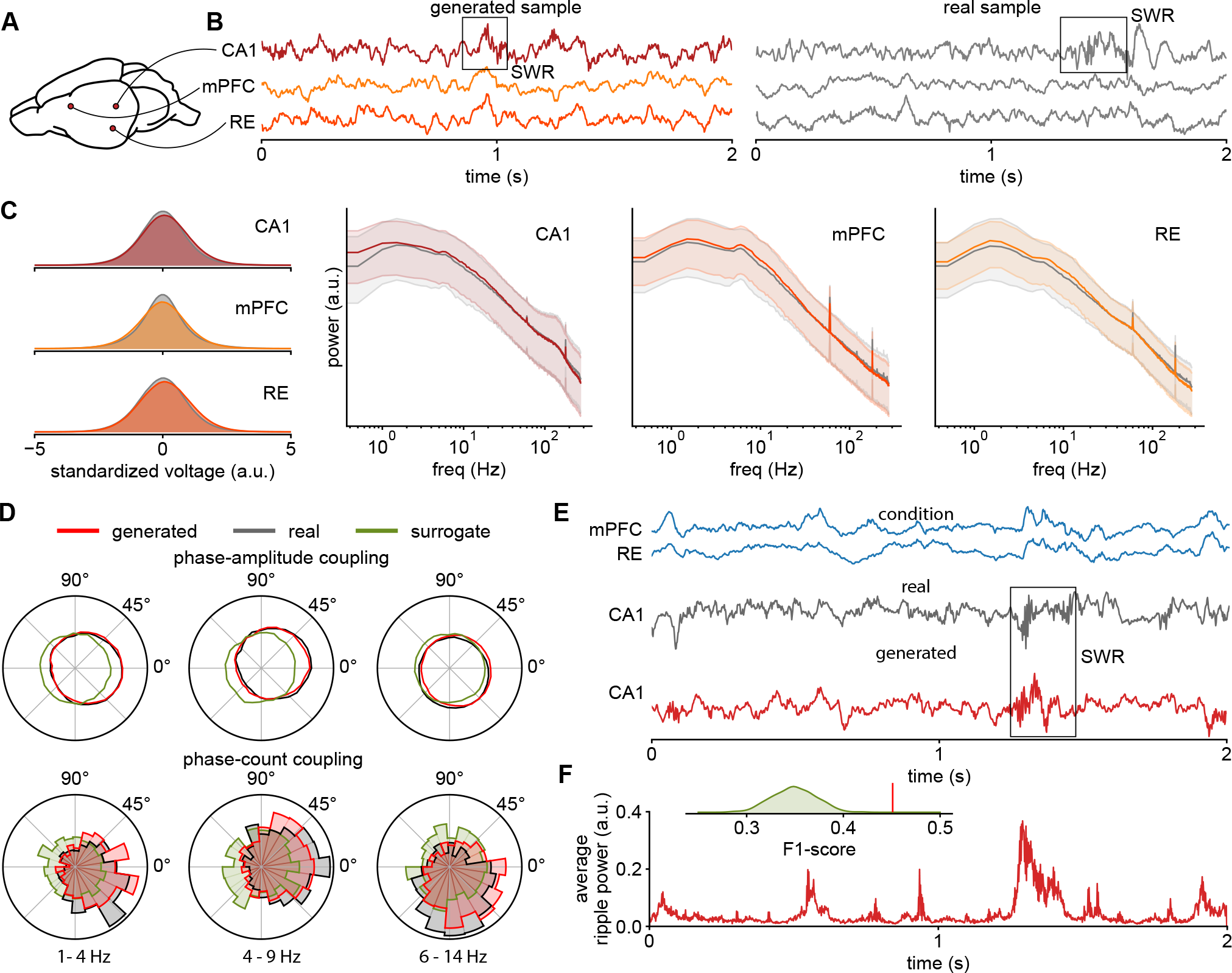
DDPM captures rat cortical-hippocampal LFP dynamics. **(A)** Schematic of rat brain showing recording locations: CA1, mPFC, and RE. **(B)** Example of time series generated by our model (left) and a real sample (right). Both examples contain sharp wave-ripples. **(C)** Marginal distributions over the standardized voltage of each channel for both real and generated data (left), as well as median and 10% / 90% percentiles of real and generated power spectra for each channel (right). **(D)** Phase-amplitude and phase-count coupling of CA1 ripples for different driver frequencies in the mPFC for real, surrogate, and DDPM-generated data. **(E)** Example of conditionally generated CA1 trace given real mPFC and RE traces. The real and conditionally generated traces contain a ripple in the same position. **(F)** Average power in the SWR frequency band (100–275 Hz) over 100 samples from the model (conditioned on the same mPFC and RE traces as in (E)) has a large peak at the same position, indicating that many SWR are generated at this location. Inset: Across the evaluation dataset, the prediction performance (“is an SWR present in both the real and predicted time series?”) measured in terms of F1 score is significantly different from the distribution of scores for randomly permuted predictions.

To assess whether our DDPM can reproduce more complex cross-regional and ripple-specific features in the real data, we measure the degree of phase-amplitude coupling (PAC) between the amplitude of CA1 ripple band (100–275 Hz) and the phase of delta (1–4 Hz), theta (4–9 Hz), and spindle (6–14 Hz) frequencies in the mPFC. We additionally isolate this analysis to the actual occurrences of ripples by thresholding the ripple band power and computing a “phase count coupling” (PCC) histogram between detected CA1 ripples and the instantaneous phase of the aforementioned driver frequencies in the mPFC (similar to Varela and Wilson (2020)). Overall, we see that ripples in both generated and real data have remarkably similar phase preferences, and for all three frequencies in the mPFC, peaking between -45 to 45 degrees (Fig. 2D). As a baseline comparison, we also created channel-wise surrogates (Venema et al., 2006) that preserve the single-channel PSD, but do not show any preferred phase coupling.

While the above demonstrates that generated samples reproduce the correct temporal relationships between CA1 ripples and slower fluctuations in the mPFC, we further investigate whether SWRs are correctly captured by our model by conditionally generating (i.e., imputing) CA1 traces given real mPFC and RE traces for a set of held-out evaluation time series. In other words, when provided with the other channels as context, can our model correctly infer when a ripple would have occurred in CA1? We use a technique known as “inpainting” to impute the CA1 channel (Lugmayr et al., 2022). This technique allows DDPMs to fill in missing values by guiding the diffusion process along observed values in the other dimensions (i.e., channels). To evaluate performance for a given (held-out) evaluation sample, we check if both the real and the imputed CA1 traces contain a ripple by thresholding ripple band power, and compute the F1 score for this classification task across the whole evaluation set. To test whether our model is better than chance at generating CA1 ripples given the appropriate context, we perform a permutation test by shuffling the binary labels of the real CA1 data (i.e., “was there a ripple or not?”) and recomputing the F1 score (Ojala and Garriga, 2010).

We provide an example of successfully imputed ripple in Fig. 2E, where the generated CA1 trace contains a ripple at exactly the same time as when the real ripple occurred around the 800ms mark. Repeating the imputation procedure 100 times on the same mPFC and RE traces, we see that the average ripple band power in the imputed CA1 traces is highest at the time that coincides with when the real ripple occurred (Fig. 2F). Across all evaluation samples, our model conditionally generates ripples significantly better than chance (F1 = 0.45, *p*_*perm*_ *<* 0.001, Fig. 2F inset). Thus, our model successfully captures within-channel temporal characteristics, as well as cross-frequency and cross-channel relationships observed in rat LFP during SWR occurrences, and can therefore be used for probabilistic generation or simulation of highly complex brain recordings. This ability to produce high-quality simulations also extends to more high-dimensional examples such as the 56-channel EEG data or the 128-channel macaque ECoG data (see Appendix).

### 2.4 DDPM-generated imputations improve neural decoding with missing data

In our second experiment, we use the AJILE12 dataset (Peterson et al., 2022), which consists of electrocorticography (ECoG) recordings from twelve human participants during naturalistic behavior while undergoing epilepsy monitoring (electrode coverage of one example participant in Fig. 3A). Using an artificially induced missing data scenario (following a similar setup as in Talukder et al. (2022)), we demonstrate how diffusion-based imputation can subsequently improve classifier performance in a neural decoding task.

**Figure 3:**
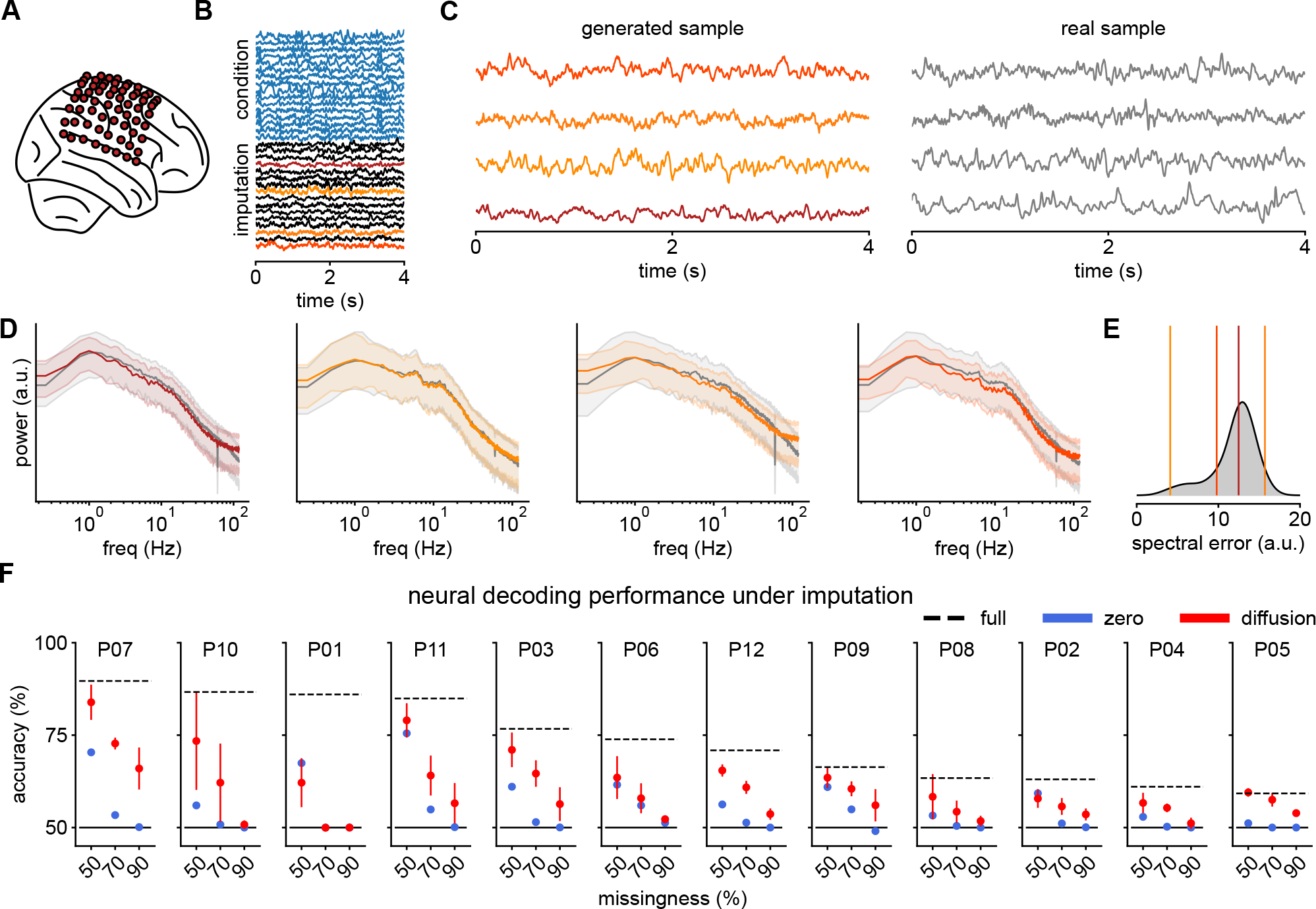
DDPM-generated imputations of human ECoG recordings improve BCI decoding. **(A)** Electrode layout of participant P07 from the AJILE12 dataset. **(B)** Example imputation of a 4-second window from P07. The first 32 channels are used as conditioning information (blue), while the remaining 32 channels are imputed (conditionally generated); four are highlighted and shown in (C). **(C)** Four randomly selected channels from the imputation (left) together with the corresponding real sample (right). **(D)** Median and 10% / 90% percentiles of real and generated power spectra for the four channels together, and **(E)** the distribution of spectral errors over all imputed channels. **(F)** Test performance of the neural decoding model for each of the 12 participants, with fully observed data (dashed), as well as with zero-imputation (blue) and DDPM-based imputation (red) under different amounts of missingness. Participants are sorted by the decoder performance on fully observed data.

For each participant, we first train a diffusion model on all channels from their first six days of recording, with the number of 4-second long training time windows varying between 148 to 1512 per participant. Unlike the model used for rat LFP data, the diffusion model for an AJILE participant is conditionally trained by randomly masking out channels and conditioning on the remaining channels to perform imputation. This conditional mask-based training can further improve imputation performance over the previously used unconditional training without masks (Tashiro et al., 2021). Details on the dataset, network architecture, and conditional versus unconditional training in Methods (Section 4).

After training the model, we test its ability to impute “missing” channels of held-out evaluation samples from the seventh day of recording by generating synthetic data for half the channels while conditioning on real data from the other half (i.e., 50% missingness, Fig. 3B). Conditionally generated synthetic traces look visually similar to real data in the corresponding channels (Fig. 3C, 4 highlighted channels), while synthetic PSDs closely match that of the real data for the imputed channels across all evaluation time series (Fig. 3D), demonstrating that DDPMs can be used for participant-specific imputation even with a substantial amount of channels missing. For the AJILE12 dataset, the quality of imputed traces is significantly improved by using OU processes instead of white noise (experiments on the effect of using the OU process versus white noise in the Appendix).

To demonstrate the utility of diffusion-based imputation, we use a binary neural decoding task of classifying the 4-second long time windows that were recorded under rest from those recorded during movement. We use per-participant random forest classifiers that were also trained on the first six days of each participant’s recording as decoders (similar to Peterson et al. (2021)), and evaluate classification performance on fully-observed evaluation windows to establish a performance upper bound (Fig. 3F, dashed). To evaluate imputation quality, for each participant, 50%, 70%, and 90% of the channels are dropped out at random from all windows of the evaluation set, and then probabilistically imputed using the trained diffusion model (as in Fig. 3B) before classifying the imputed time series using the same random forest classifier. We repeat this random channel dropping procedure 5 times, and report the average and standard deviation in Fig. 3F (red). We also compare diffusion-based imputation to a zero-imputation baseline (Fig. 3F, blue)—a strategy often employed in such neural decoding paradigms when faced with missing data (Talukder et al., 2022).

Across 12 participants and 3 missingness levels, we observe that diffusion-based imputation leads to better classification performance than zero-imputation in 27 out of 36 cases. Remarkably, for some participants (e.g., P05, P07, P09, Fig. 3F), diffusion-based imputations almost recover the original decoding performance using all channels. Overall, classification performance after imputation appears highly participant-dependent. Nevertheless, diffusion-based imputation significantly improves neural decoding accuracy compared to zero-imputation, even with a significant proportion of channels missing, while achieving similar or better performance compared to more tailored imputation methods for neurophysiological recordings (e.g., Talukder et al. (2022)).

### 2.5 Class-conditional DDPMs can generate and evaluate brain state-dependent recordings

In our third experiment, we use DDPM to model whole-brain surface electrocorticography (ECoG) from non-human primate (macaque monkey) sampled at 1000 Hz (Yanagawa et al., 2013). In this experiment, we model 12 randomly selected channels (out of 128, Fig. 4A) during two distinct behavioral states: awake, a control condition where the animal is head- and arm-restrained but otherwise freely behaving, and anesthetized, where the animal is fully unconscious following the induction of general anesthesia. Our class-conditional diffusion model can accurately generate synthetic recordings under both states, which can be used to train a classifier without giving it access to real data. In addition, we can evaluate likelihoods under the learned generative model, which can be used for state classification and outlier detection of real recordings.

**Figure 4:**
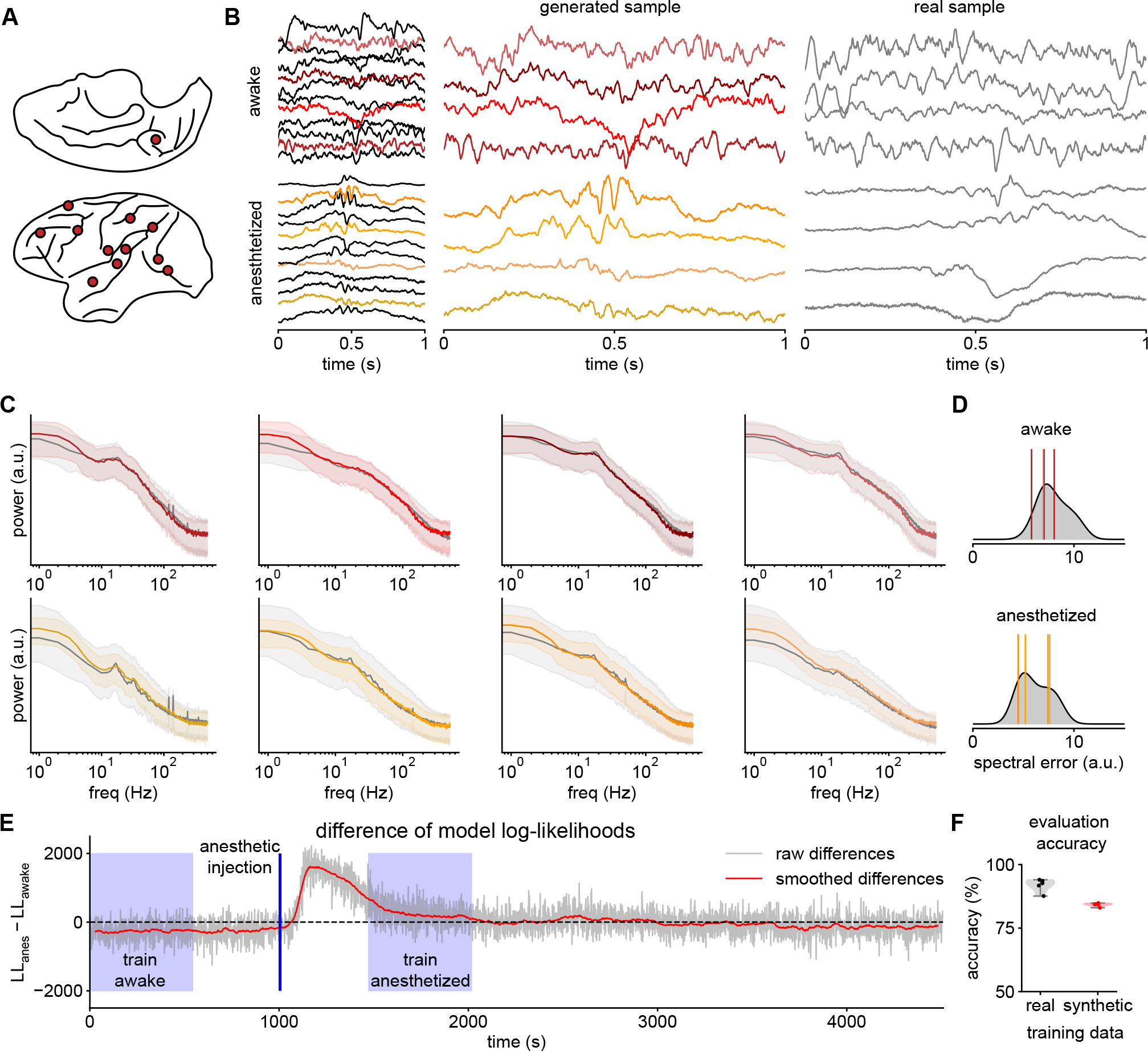
Conditional DDPM learns brain state-dependent macaque ECoG recordings. **(A)** Electrode locations in the macaque brain of the 12 randomly selected channels used for modeling. **(B)** Examples of conditionally generated time series for both the awake and anesthetized condition. 4 randomly selected channels are shown in more detail (left) together with real samples (right). **(C)** Median and 10% / 90% percentiles of real and generated PSDs for the four channels (left) stratified by condition, together with **(D)** the corresponding distribution spectral errors of all 12 channels (right). **(E)** Difference of log-likelihoods over 1.5 hours of recording. A positive difference indicates a classification towards “anesthetized”, a negative difference towards “awake”. Periods used for model training are shaded in blue. **(F)** Real-data test accuracy of a brain-state classifier trained on real (black) vs. DDPM-generated synthetic data (red).

After training our model on a total of 870 1-second long windows from both the awake and anesthetized conditions, we conditionally generate the same number of synthetic time series for evaluation. For both the awake and anesthetized condition, the real and generated example time series are visually indistinguishable from each other (Fig. 4B, 4 out of 12 channels are highlighted). Notably, synthetic data during the awake state contains more higher frequency fluctuations and oscillations lasting several cycles, while abrupt lower frequencies fluctuations dominate during the anesthetized state. Across all 12 modeled channels, the PSDs of real and synthetic data closely match in both conditions (Fig. 4C, D).

In addition to generating synthetic recordings in a state-dependent manner, class-conditional DDPMs can also evaluate the likelihoods of recordings under each condition, i.e., how likely is it to observe a given recording under awake vs. anesthetized states? Using the difference of log-likelihoods from our trained models to classify previously unseen evaluation time windows from either state, we achieve an evaluation accuracy of 83% on the balanced classification task. Additionally, when we evaluate likelihoods over a 1.5-hour segment of the recording in a sliding-window manner, we observe that the difference in log-likelihood between anesthetized and awake conditions is negative to start, and sharply increases after the induction of general anesthesia—even though the diffusion model does not have access to this information, nor was it trained on nearby segments of the recording (Fig. 4D, training segments in blue). As time elapses, the difference of log-likelihoods slowly decreases towards the “awake level” (negative) as the anesthesia wears off, demonstrating how such generative models can be applied for continuous state prediction.

Additionally, unlike with purely discriminative classifiers, the ability to evaluate likelihoods allows us to use our generative model to perform outlier detection, which we demonstrate through two experiments. In the first experiment, for all time series in the evaluation set, we replace half of the channels with white noise to simulate complete signal loss in some of the recording electrodes. In the second experiment, we instead reverse half of the channels time-wise, which is unlikely to happen in the real world, but creates time series that are visually and statistically similar to the real data, thus representing a difficult out-of-distribution test scenario. Using the likelihood of the real and outlier data provided by the diffusion model as a score for classification, the AUC for time-wise flipped outliers is 0.63, while the AUC for white noise outliers is 1.0. When comparing pairs of time-wise flipped outliers with their corresponding real time series, the likelihood of the real time series is larger in 98% of the cases, demonstrating the model’s ability to distinguish data that have the same marginal distribution per channel, but are nevertheless out-of-distribution.

Finally, to further demonstrate the utility of using diffusion models to generate realistic neurophysiological recordings, we perform brain state classification based on synthetic data. In many situations, the original data is sensitive due to privacy concerns, e.g., when collected as a part of medical examination, and therefore cannot be publicly shared, but may otherwise be useful for applications such as training of biometric algorithms. In such situations, synthetic recordings that preserve specific aspects of the recording while anonymizing others may be valuable. Here, we use the trained DDPM to generate synthetic data from awake and anesthetized conditions, and train a neural network-based discriminative classifier (as described in Wang et al. (2017)) solely on this synthetic data. The trained classifier is then evaluated on a held-out evaluation set of real recordings, while never having seen real data before. Since the diffusion model captures the conditional distribution of the two classes well, the classifier trained on synthetic data achieves an accuracy of 84% when classifying real data (Fig. 4E), which is close to the accuracy of a classifier trained on real data (92%). We thus demonstrate the ability to create artificial datasets that can be shared to facilitate further research with reduced privacy concerns.

## 3 Discussion

### 3.1 Summary

Our experiments show that denoising diffusion probabilistic models (DDPMs), together with a flexible convolutional denoising network and the use of the Ornstein-Uhlenbeck processes, are a powerful tool that can accurately model highly multivariate and densely sampled neurophysiological data, providing features, and more importantly, realistic synthetic data, useful for applications relevant to neuroscience. The power of DDPMs lies in their generality, as they provide a general-purpose generative model that can be used for many different applications ranging from unconditional (i.e., simulation) and conditional generation, to imputation and likelihood evaluation.

We demonstrate these applications here on three different datasets of neurophysiological recordings with channel counts ranging from 3 to 128 channels, from a variety of brain regions, recording modalities, and species. Overall, our models accurately capture data distributions in time and frequency domains, as well as detailed cross-channel and cross-frequency dependencies like sharp wave-ripples. On an imputation task, they achieve a performance that is on par with or better than tailored imputation methods for neurophysiological recordings. Finally, we show successful applications of class conditional models in classification, outlier detection, and synthetic training data generation.

These features make DDPMs a viable tool for neuroscientists and clinicians studying the complex dynamics embedded in neurophysiological recordings. After successful training, DDPMs can be used to generate unlimited amounts of data as input to neuroscience simulators. When neurophysiological data are subject to strict privacy requirements, as may be the case with human clinical data, neuroscientists and clinicians can use DDPMs to generate artificial datasets that can be shared with collaborators with less concern. In addition, the accurate imputations provided by DDPMs can eliminate the need to discard large amounts of data or to modify an already trained neural decoder. Finally, DDPMs output log-likelihoods that provide users with a model to perform simple classification tasks or outlier detection at no additional training cost.

### 3.2 Other potential applications

Trained DDPMs can be used for a variety of other applications that we do not show in this work, such as training data augmentation, as well as imputation of more complex missingness patterns, which includes the task of time series forecasting.

Instead of completely replacing training datasets with artificial ones, as described in the data privacy use case (Fig. 4F), it is possible to augment existing real training data with synthetic data generated by a DDPM to improve the performance of another model (e.g., a classifier). This additional training data could lead to performance improvements, especially if the relevant model is difficult to regularize and prone to overfitting. DDPMs can provide tailored “noise”-augmented data samples that capture complex relationships other surrogate methods may not. Throughout our experiments, the DDPMs were robust to overfitting and were able to achieve good sample quality with only a few hundred training time windows of one to four seconds long. In addition, DDPMs can be trained on a separate, larger dataset to generate synthetic “pre-training” data for a classifier model that is later fine-tuned on task-specific but smaller datasets, i.e., to facilitate transfer learning.

DDPMs can also handle more complex missingness patterns. In this work, we have demonstrated the ability of DDPMs to perform cross-channel imputation, although completely missing channels represent only one “pattern” of missingness that can occur in real neurophysiological recordings (i.e., dead electrode). Many other patterns of missingness are possible, such as when some time points within a recorded channel are missing due to momentary artifacts. Imputation in this context would mean filling in these missing measurements given information from other channels *and* observed values in the channel itself. These more complex missingness patterns can be realized with diffusion models under both training strategies: For unconditional training without masks, all possible missingness patterns work out of the box via “inpainting” (i.e., Fig. 2E). For conditional mask-based training, the training strategy needs to be generalized from dropping out whole channels to more complex missingness patterns. Tashiro et al. (2021) discuss several strategies for this purpose. A special case is the common task of time series forecasting, where the “missing” values are simply unobserved time points in the future. Like imputation, time series forecasting can then be realized by diffusion models trained unconditionally without masks or conditionally with masks.

An additional benefit of our fully convolutional denoising network is that it allows users to generate neurophysiological recordings of arbitrary length, despite training on fixed-size time series. To generate long synthetic recordings, the denoising network is simply applied to the entire time series without any further technical modifications (though dependencies longer than the timescale of training segment may not be correctly modeled). We have not used this capability here, but it will be important when generating input to a simulation where long synthetic recordings are required.

### 3.3 Limitations

Although diffusion models are a very popular type of deep generative models today, they are not without limitations. Because DDPMs generate samples by successively denoising them over (typically) hundreds of denoising steps, inference is computationally intensive, especially when compared to other families of deep generative models such as VAEs or GANs. Much recent work has focused on speeding up the inference time of diffusion models, often by tolerating a small degradation in sample quality (Song et al., 2021a, Karras et al., 2022, Lu et al., 2022). While in this work we applied the standard formulation of DDPMs without further speedup, all progress in this direction is directly applicable to our setting and provides a potential remedy for the comparatively long inference times.

Furthermore, diffusion models typically operate on the dimensionality of the original data space. In neuroscience, however, there has been much interest in lower-dimensional state-space representations of high-dimensional neurophysiological recordings, especially of neuronal population spiking data (Vyas et al., 2020), but more recently also of continuous time series such as LFPs (Gallego-Carracedo et al., 2022). One way to obtain such low-dimensional neural representations with generative models is to use VAEs (e.g., like in Pandarinath et al. (2018)), which model the original data distribution via a mapping to a lower-dimensional space of fixed dimensionality. This encoding property is not automatically fulfilled by DDPMs, but combinations of encoder-decoder architectures with a diffusion model in the latent space have been successfully applied to image generation and fMRI decoding (Rombach et al., 2022, Chen et al., 2023). Similar approaches are possible for neurophysiological recordings and may provide low-dimensional representations, along with increased computational efficiency. In particular, a shared latent space could be beneficial in applications similar to our imputation experiment on the AJILE12 dataset (Fig. 3). Unlike the autoencoder-based model described in Talukder et al. (2022), we trained an individual DDPM for each of the twelve participants. Therefore, a potential extension of our current model is to jointly train a latent diffusion model using participant-specific encoder and decoder layers, which could increase performance and robustness due to its ability to generalize over different participants.

As discussed earlier, synthetic training data can alleviate privacy concerns. However, our DDPM-generated synthetic data is private in the sense that the minimum difference between time series in the real training data is similar to the minimum difference between time series in the real and generated data. While intuitive, this measure does not provide theoretical guarantees against any form of reverse engineering or memorization (van den Burg and Williams, 2021).

Lastly, a persisting challenge in modeling neurophysiological recordings is to capture very long-range dependencies over timescales of minutes or even hours. In our training setup, the time series extracted from a long neurophysiological recording are treated as independent samples by the model. Thus, temporal dependencies beyond the length of the time series used for training are not captured. Naturally, for any given dataset, long-range and cross-scale dependencies are much harder to capture due to data sparsity. Thus, building deep generative models that are more data efficient and can capture long-range dependencies by training on only a few samples is a challenging but exciting avenue for further research.

### 3.4 Conclusion

Together with a flexible convolutional denoising network and the use of the Ornstein-Uhlenbeck processes, DDPMs provide a powerful and flexible type of generative model for neurophysiological recordings that can generate high-quality samples, and can be used in a variety of applications relevant to neuroscience like simulation, imputation, or brain-state classification.

## 4 Methods

### 4.1 Denoising Diffusion Probabilistic Models

To train our diffusion models, we consider a training dataset of *N* time windows extracted from neurophysiological recordings 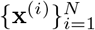 where **x**^(*i*)^ ∈ R^*C×L*^ with *C* channels of fixed length *L* time steps. During sampling or inference, arbitrary time series lengths are admissible due to our fully convolutional network architecture.

#### Background on Denoising Diffusion Probabilistic Models

Our primary goal is to learn a parametrized probability distribution *p*_*θ*_(**x**) on R^*C×L*^ that is close to the original data distribution *p*(**x**). *p*_*θ*_(**x**) can then be used for many different tasks like imputation, generation, or likelihood evaluation. We use denoising diffusion probabilistic models (DDPM) (Ho et al., 2020):

DDPMs work by modeling two processes, the reverse (denoising) and forward (noising) diffusion process, over a sequence of latent variables **x**_*t*_, *t* = 1, …, *T*, where *T* is a pre-specified process length. Note that *T* is the number of diffusion steps, a hyperparameter in the generative model, and not the length of the time series being modeled (*L* above). The forward process is given by a Markov chain

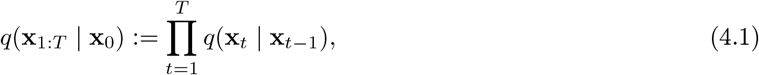

where the noising distribution

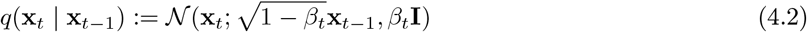

iteratively adds Gaussian noise controlled by the noise level *β*_*t*_ *>* 0, resulting in an evolution from the data distribution, *p*(**x**_0_), to the noise distribution, *p*(**x**_*T*_), over the course of *T* diffusion steps. This forward process is chosen by the user beforehand and thus not learned.

As its name suggests, the reverse process reverses the forward process and is given by

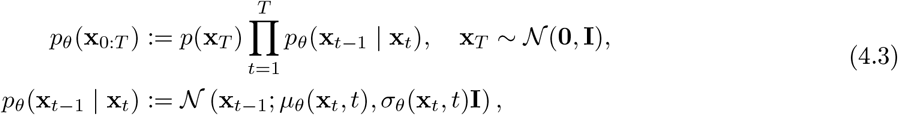

i.e., the reverse conditional distribution is also Gaussian and its mean and variance are parametrized functions of the current diffusion step *t* and the noised data point **x**_*t*_, and therefore need to be learned. DDPMs parameterize *p*_*θ*_(**x**_*t−*1_ | **x**_*t*_) as a denoising distribution,

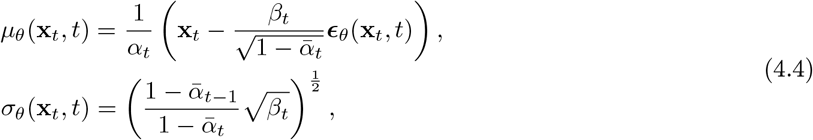

where ***ϵ***_*θ*_ is a neural network with learnable weights *θ*.

In other words, the neural network ***ϵ***_*θ*_(**x**_*t*_, *t*, cond) acts as “denoiser” that, together with the diffusion time step *t* and, optionally, additional conditioning information cond, produces a denoised version of the data point (in this case a time series) as the basis for the next denoising step.

The denoising network can be optimized by computing a variational lower bound on *p*_*θ*_(**x**_*t*_ **x**_*t−*1_), which results in the full (simplified) training objective:

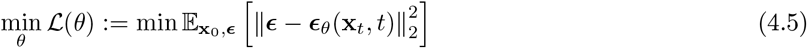

with 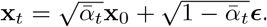

#### Denoising Diffusion Probabilistic Models for neurophysiological time series

For our application of modeling neurophysiological time series, we extend the standard DDPM setting with two modifications:

1. First, we use a flexible convolutional neural network architecture based on structured convolutions as our denoising network (Li et al., 2023). This architecture is able to accurately model long (thousands of time points) and highly multivariate time series (over 100 channels).
2. We replace the white noise process used in the standard DDPM by general Gaussian processes. This allows us to include “prior information” about the modeled time series into the generative process (Biloš et al., 2022). In this paper, we specifically focus on the Ornstein-Uhlenbeck process due to its advantageous numerical properties and its relevance for neurophysiological recordings, in particular the inverse power law scaling over frequencies.

#### Network architecture with structured convolutions

Our denoising network consists of interleaved layers of structured convolutions (Li et al., 2023) and linear layers that “mix” information between channels. Structured long convolutions are created by concatenating several linearly interpolated and exponentially downscaled convolutional kernels. The resulting convolutional kernel can be as long as the time series itself. This construction is inspired by recent advances in deep state space models (Alcaraz and Strodthoff, 2022, Gu et al., 2022) and allows for long yet parameter-efficient convolutional kernels that can model complex and long-range dependencies within a given time series. To make the use of these large convolutional kernels tractable, the convolution operation is computed by element-wise multiplication in the Fourier domain after applying a Fast Fourier Transform. This approach drastically reduces the computation time for large kernels and long time series.

The input layer is a standard convolutional layer (LeCun et al., 1995) that maps the input as well as the diffusion time embedding and other conditional information into a high-dimensional latent space. We parameterize the size of the latent space per channel, i.e., we specify the number of latent dimensions per channel. The full dimensionality of the latent space is then given by the number of channels times the latent dimension per channel. The output layer is also a standard convolutional layer that maps the hidden activations to the desired output dimensionality. The hidden layers consist of blocks of structured convolutions followed by linear layers. The linear layers “mix” the output from the structured convolutions, which operate separately on each hidden channel. Unlike the original implementation of structured convolutions, we introduce the number of scales as a new hyperparameter. This means that the overall size of the structured convolutional kernel is given by kernel_size 2^num_scales*−*1^.

In our experiments, especially when training on neurophysiological recordings with high channel count, we noticed that fully parametrized linear mixing layers can cause convergence issues. The number of weights in the linear layers grows quadratically with the full latent dimensions and can quickly dwarf the number weights in convolutional layers. To avoid this overparameterization and alleviate the convergence issues, we sparsify the weight matrices of the linear layers. To do so, we decrease the number of active weights in off diagonal blocks (see Appendix Fig. 5). The size of the off diagonal blocks is a hyperparameter.

**Figure 5:**
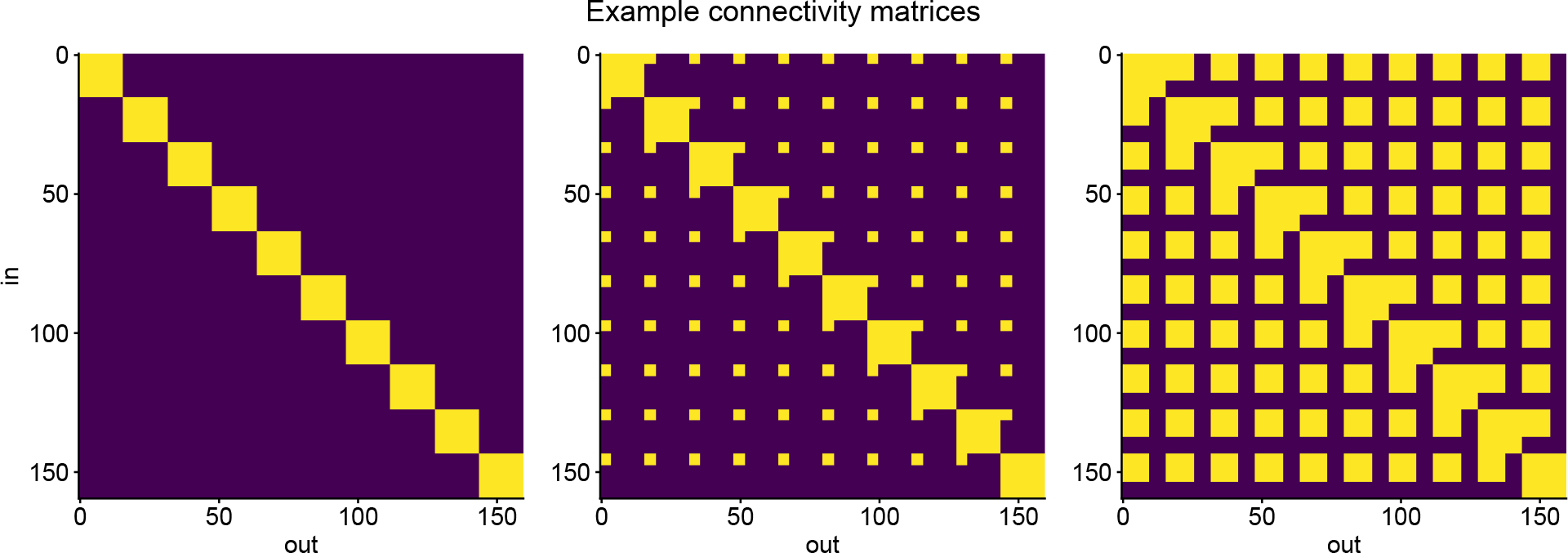
The linear “mixing” layers that follow the structured convolutions are parametrized as block diagonal matrices. To avoid overparameterization of the network for neurophysiological recordings with very large channel counts, we sparsify the corresponding matrices by reducing the size of the off-diagonal blocks. Three examples with increasing off-diagonal block size are shown. Yellow indicates that the weight is part of the network, blue indicates no connection.

Throughout our architecture, we use GELU activation functions (Hendrycks and Gimpel, 2016) and batch normalization (Ioffe and Szegedy, 2015). Importantly, our architecture does not perform any compression in time. This allows the denoising architecture to operate on time series of arbitrary length.

#### Alternative noise processes as diffusion process

It is straightforward to replace the white noise process commonly used in DDPMs with other Gaussian processes (Biloš et al., 2022). Instead of using white noise from 𝒩(**0**, *I*), the (forward) diffusion process is performed with noise sampled from a general Gaussian process 𝒩(**0**, Σ) with prespecified covariance Σ. The objective from Eq. (4_5) is also changed to include Σ:

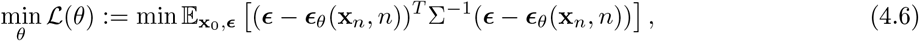

where 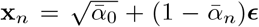 and ***ϵ*** (**0**, Σ). This replacement is useful for providing the model with domain knowledge about the recordings and can lead to better empirical performance.

One such example of domain knowledge in the context of neurophysiological recordings is continuity. Neurophysiological recordings are typically continuous and do not exhibit large and abrupt jumps. In addition, for many neurophysiological recordings, the frequency spectrum exhibits inverse power law scaling (Pritchard, 1992).

In this paper, we use the Ornstein-Uhlenbeck (OU) process, which is a continuous noise process with power-law frequency spectrum to encode this domain knowledge. The OU process is a stationary Gauss-Markov process with many applications in physics and financial mathematics (Chan et al., 1992). As the “roughest” member of the family of Matern kernels, it is continuous, but not differentiable.

Using the OU process is computationally feasible, even for long time series. While in general sampling from Gaussian Processes scales cubicly with the length of the time series, for the OU process it is possible in linear time. This is because the OU process can be described equivalently by the stochastic differential equation,

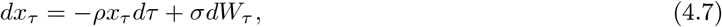

where *W*_*τ*_ is the Wiener process. Furthermore, since the OU process is a stationary Gauss-Markov process, its precision matrix is banded and the modified DDPM training objective can also be computed in linear time. For both sampling and computing the training objective, we apply the OU process independently for each time series channel. However, we find that this modification does not improve sample quality in all datasets. See experiments on the effect of using the OU process versus white noise in the Appendix.

### 4.2 Model training and inference

Our convolutional denoising networks are trained with stochastic gradient descent on the objective given in Eq. (4_6). We train all our models using the AdamW optimizer (Loshchilov and Hutter, 2019) and choose hyperparameters by inspecting the quality of the generated data with respect to the training set. See the Appendix for all network and training hyperparameters. To sample and impute using the trained denoising network, a sample from the Gaussian distribution (white noise or OU) is gradually denoised following the standard DDPM denoising procedure (Ho et al., 2020).

DDPMs naturally lend themselves to imputation by guiding the diffusion process based on observed data points (Ho et al., 2020). For image-based diffusion models this process is usually known as “inpainting” (Lugmayr et al., 2022). It works by replacing the observed part of the image—or time series—with the analytical latent state of the forward diffusion process after each denoising step. This way, the reverse diffusion process is “guided” along the observed part of the time series. However, this approach does not provide the correct conditional distributions of the form *p*(**x**_observed_ | **x**_missing_), which can hurt imputation performance (Tashiro et al., 2021).

To alleviate this issue, it is possible to train conditional diffusion models directly: Here, the condition is given as an input mask to the denoising network to compute the correct conditional distribution. To account for different patterns of missingness at imputation time, many condition input masks need to be sampled during training. In this work, we use the random sampling strategy (Tashiro et al., 2021) and randomly drop out channels for a given time series following a two step process: First, we uniformly sample the number of channels to drop, and then uniformly drop out channels according to this number. The denoising network is then conditioned on the remaining channels and the training objective is computed. We use this masked-based conditional training only for the AJILE12 dataset. For the other two datasets, the models are trained without masks. All models are implemented and trained with the PyTorch library (Paszke et al., 2019).

### 4.3 Datasets

We perform our three experiments on three different datasets of publicly available neurophysiological recordings.

The dataset for our first experiment (Fig. 2) is a three-electrode LFP recording from rats recorded at 600 Hz at the medial prefrontal cortex (mPFC), hippocampal CA1 region, and the thalamic nucleus reuniens (RE) (Varela and Wilson, 2020)^1^. It consists of a total of 3.5 hours of recordings from three different rats. During the recording, rats were freely behaving with the possibility to rest and underwent cycles of sleep and wakefulness. For our experiment, we used the LFP data from the first and third recording (52 and 93 minutes).

The AJILE12 dataset used for our second experiment (Fig. 3) consists of human ECoG data recorded during naturalistic behavior from 12 different participants undergoing epilepsy monitoring (Peterson et al., 2022)^2^. It was recorded at 500 Hz over several days, and a total of 1280 hours. The length of recording per participant ranges from 70 to 120 hours. The number of electrodes per participant ranges from 64 to 126. In addition to the neurophysiological recordings, a variety of behavioral and movement event related metadata are provided.

The dataset for our third experiment (Fig. 4) consists of ECoG recordings from non-human primates (macaque monkey) recorded at 1000 Hz via surface 128 electrodes placed inside the cranium, directly on top of the cortex (Yanagawa et al., 2013)^3^. We use data from only one macaque (Chibi). During the recording, the macaques were injected with an anesthetic (propofol) and different neurophysiological states were annotated. We use the data from the anesthetized and awake-eyes-closed condition (73 min of recording).

For our initial example (Fig. 1E-H), we use data from participant P02 of the BCI Challenge @ NER 2015 (Margaux et al., 2012)^4^. The data was recorded at 200Hz with 56 passive EEG sensors placed with the extended 10-20 system. The participants were tested under the “P-300 Speller” paradigm, where words are spelled out letter-by-letter by flashing screen items and measuring the evoked response (Farwell and Donchin, 1988).

### 4.4 Data preprocessing

For our datasets of rat LFP and macaque ECoG recordings, we perform channel-by-channel normalization by standardizing the recording over its entire duration. After normalization, we create the set of time windows that are used to train and evaluate the model. For the rat LFP recordings, we took 2-second time windows consisting of 1200 time points resulting in a total of 4350 time series. Similarly, for the macaque ECoG dataset, we took 1-second time windows consisting of 1000 time points resulting in a total of 1088 time series. After creating the time series, we then randomly divide them into a training and an evaluation set (80% training, 20% evaluation).

The data of each participant in the AJILE12 dataset consists of seven days of recording. We follow the setup and preprocessing described in Peterson et al. (2021): The ECoG traces are downsampled before extracting time windows corresponding to either movement or rest. For each participant, the final dataset consists of 4-second time windows with a sampling frequency of 250 Hz with an equal amount of time series recorded under rest and movement (i.e., class-balanced). Again, we normalize the time series channel-by-channel by aggregating over all time series in the training set. The normalization of the training set, which consists of the first six days of recording, is applied to the evaluation set, i.e., the seventh day of recording. Furthermore, extreme outlier channels, where the difference between the minimum and maximum value is five or more times the interquartile range, are detected and masked during training and evaluation. The number of recorded channels varies widely between participants, ranging from 64 to 126. Similarly, the number of training and evaluation time series varies considerably between participants. However, the network hyperparameters are shared between participants and no attempt was made to fine-tune the imputation for an individual participant.

Unlike the DDPM, the random forest neural decoder is trained and evaluated on the unnormalized AJILE12 data (see Peterson et al. (2021) for details). Imputations are obtained by imputing normalized time series and then reversing the normalization for both the observed and imputed channels using the normalization constants from the training data.

For our EEG dataset, we extract 1.3 seconds of the recording immediately following a presented stimulus. Since we only use data from participant P02, this results in 340 time series (i.e., trials) consisting of 260 time points each. After extraction, we band-pass filter the time series with a frequency window between 1 and 40 Hz using a 5th order Butterworth filter. We then again perform channel-by-channel normalization by aggregating over the time series.

### 4.5 Analyses and Metrics

#### Power spectral densities and spectral error

Throughout the experiments, we compare the distributions of (cross-) power spectral densities (PSDs) between channels of real and generated data. PSDs are computed with a Fourier transform over the whole the time series. For a given one-dimensional time series *x* and frequency *f*, the PSD is denoted by *S*_*xx*_(*f*). Since we are always interested in capturing the full distribution of PSDs over a set of time series, we compute the point-wise median and percentiles: Here, point-wise means that, for a set of real or generated time series 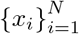, the median median[*S*_*xx*_(*f*)] over all PSDs is computed at each frequency *f*, and similarly for the 10% and 90% percentiles.

To quantify how well the distribution of PSDs of the real and generated data match in a single metric, we compute the spectral error in log space. The spectral error is defined as

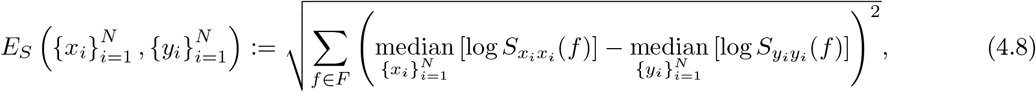

where *f* ∈*F* are frequency bins and median[log *S*_*xx*_(*f*)] is the median log-power at a fixed frequency *f* over all generated (*y*_*i*_) or real (*x*_*i*_) time series. All computations related to the power spectrum are performed with the numpy (Harris et al., 2020) and Scipy (Virtanen et al., 2020) libraries.

#### Sharp wave-ripple detection and cross-channel couplings

For our experiments with the rat LFP data, we detect ripples in the CA1 channel. Following the heuristic approach of Varela and Wilson (2020), we perform this detection by band-pass filtering both real and generated CA1 traces within the ripple frequency band from 100 to 275 Hz using a finite impulse response (FIR) filter. Ripples are detected when the power of the time series is more than three standard deviations greater than the mean. The mean and standard deviation of the filtered power are calculated over the entire set of real time series. Crucially, to ensure a fair comparison between a set of real and a set of generated CA1 traces, the mean and standard deviation from the real data are used to detect SWRs in the generated data. SWRs with less than 10 time points (17ms) are fused. After fusing, a SWR must have a minimum length of 20 time points (33ms). Detected ripples below this threshold are discarded.

In our SWR prediction experiment, we impute the CA1 channel given the other two channels and detect SWRs in the real and imputed channel for all time windows in the evaluation set. If there are no SWRs or at least one SWR in both the real and generated channel, the prediction is counted as correct. We then compute the F1 score based on the number of correct and incorrect predictions. To test if the prediction performance is significantly better than random, we shuffle the set of generated time series and evaluate the F1 score. This procedure is repeated a 1000 times to compute the permutation test statistic (Ojala and Garriga, 2010).

We also use detected SWRs to compute a “phase-occurrence coupling” against the phase of delta (1–4 Hz), theta (4–9 Hz), and spindle (6–14 Hz) frequencies in the mPFC. The mPFC phase is calculated using the Hilbert transform after band-pass filtering (FIR, Hamming) with the corresponding frequency band. Given the mPFC phase, the time point with maximal power within a detected SWR is used to extract the phase of the SWR occurrence, and the counts are binned into 21 equally spaced phase-bins (from -180 to 180 degrees) to create the histograms (Fig. 2D).

Finally, we compute the phase-amplitude coupling (PAC) between the CA1 amplitude in the ripple band (100–275 Hz) and the mPFC phase of the previously mentioned driver frequencies. Phase and amplitude in the respective frequency bands are calculated using the Hilbert transform, and the average amplitudes occurring at 31 equally spaced phase-bins (from -180 to 180 degrees) are reported (Fig. 2D).

#### Details on the imputation experiment and neural decoder

For our imputation experiment, we follow the setup described in Talukder et al. (2022): We impute artificially dropped channels and investigate the effect of this imputation on the performance of a neural decoder. As a baseline imputation method, we use zero imputation. The neural decoding task is to discriminate between time series recorded at rest and time series recorded during movement. The neural decoder used is a random forest, which works by “flattening” a time series into a large feature vector. We train three decoders over three training folds as in Peterson et al. (2021) using the same hyperparameters (maximum tree depth and number of estimators). No attempt was made to further optimize the random forest models. Finally, the average test performance over all training folds is calculated for all settings, i.e., fully observed, zero imputation, and DDPM-based imputation.

To evaluate the performance of DDPM-based imputation, we randomly select 50%, 70%, or 90% of the channels from a given participant. For each time series in the evaluation set, this set of channels is dropped and then imputed with zeroes or imputed using the trained DDPM. We then apply the previously trained decoder to both imputed evaluation sets and compute the evaluation accuracy. This process is repeated 5 times for each missingness level to evaluate the imputation performance over different sets of dropped channels. We report the mean and standard deviation over the 5 repetitions (Fig. 3).

#### Likelihood computation and brain-state classification

Likelihood computation in DDPMs can be realized by solving the probability flow equation (Song et al., 2021b). In our class-conditional model for the macaque ECoG data, we compute log-likelihoods for both the awake and anesthetized states. That is, given a time series **x**, we get log *p*_*θ*_(**x** *c* = awake) and log *p*_*θ*_(**x** *c* = anesthetized). These values can be used to perform classification by selecting the class with the higher log-likelihood for a given sample. We perform this classification for all time series in our class-balanced evaluation set and report the overall classification accuracy. For the time-resolved brain state classification in Fig. 4E, we simply compute this log-likelihood difference over 1-second sliding windows.

For outlier detection, we are interested in the total log-likelihood log *p*_*θ*_(**x**) of sample **x**. We compute this total log-likelihood for each evaluation time series and correspondingly generated outliers (white noise or time-wise flipped) according to the law of total probability. Then, the area under the receiver operator characteristic (AUC) of the time series against both types of outliers is computed.

For the brain state classification with synthetic data, we use the convolutional neural network classifier described in Wang et al. (2017). The classifier is trained twice, once on real data and once on an equal amount of DDPM-generated data, and then evaluated on a held-out evaluation set of real data. Both the training and evaluation datasets consist of an equal number of awake and anesthetized time windows, and we compute the evaluation accuracy to compare the performance between the real and synthetic settings. The classifier is trained using the AdamW optimizer (Loshchilov and Hutter, 2019) and binary cross entropy loss with identical network and training hyperparameters in both settings.

## 5 Acknowledgments

JV is supported by the International Max Planck Research School for Intelligent Systems (IMPRS-IS) and the AI4Med-BW graduate program. RG is supported by the European Union’s Horizon 2020 research and innovation program under the Marie Sk łodowska-Curie grant agreement No. 101030918 (AutoMIND). JHM, RG are members of the Machine Learning Cluster of Excellence, EXC number 2064/1–390727645. This work was supported the German Federal Ministry of Education and Research (BMBF): Tübingen AI Center, FKZ: 01IS18039A. We would like to thank Auguste Schulz and Matthijs Pals for feedback on the manuscript and Mackelab members for discussions throughout the project.

## 6 Appendix

### 6.1 Ornstein-Uhlenbeck process as diffusion process

In our experiments, we sometimes use the OU process instead of the classically used white noise as our diffusion process. This substitution is possible because the OU process belongs to the family of Gaussian processes. See Biloš et al. (2022) for more details.

The training and sampling schemes are shown in Algorithm 1 and Algorithm 2 and differ slightly from those given in Biloš et al. (2022). Note that the extension to general Gaussian processes has a computational overhead compared to the original training objective. During training, the computation of the Mahalanobis distance scales quadratically with the length of the input. Sampling from a Gaussian process also requires computing a Cholesky decomposition. However, this decomposition only needs to be computed once, and can also be used to compute the Mahalanobis efficiently by solving a lower triangular linear system based on the Cholesky decomposition.

#### Algorithm 1

Training

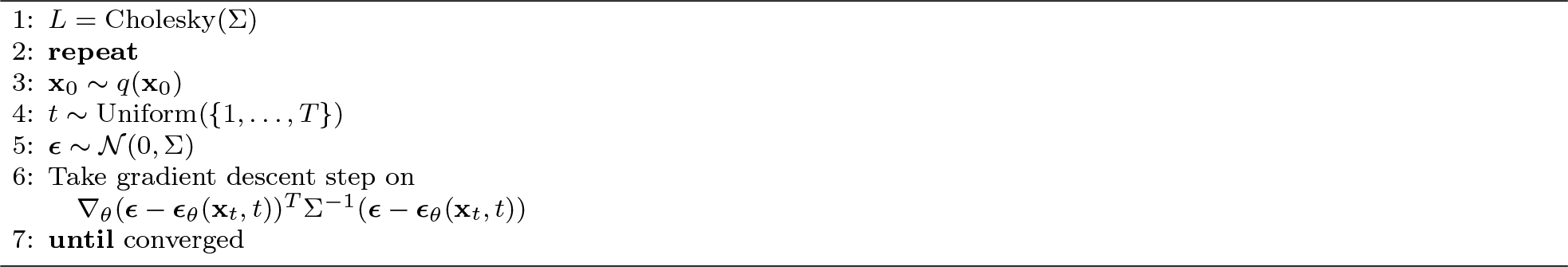

#### Algorithm 2

Sampling

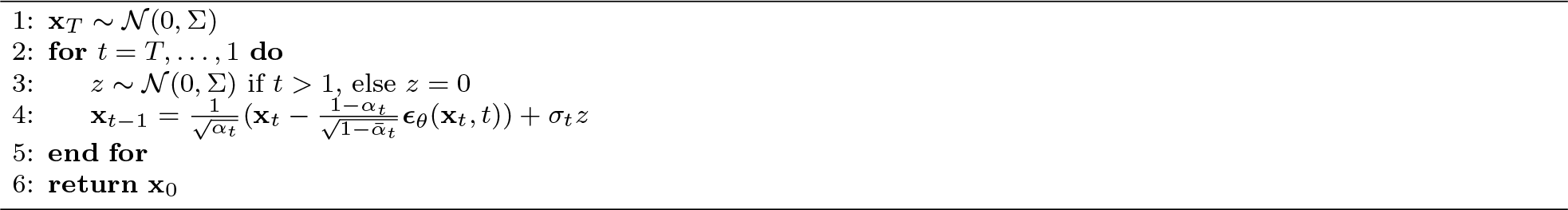

As described in the main text, the computational requirements scale more favorably for the special case of the OU process. Its formulation as a stochastic differential equation allows sampling in linear time. Furthermore, the OU process has a tridiagonal precision matrix, which allows the the Mahalanobis distance in the training objective to be computed in linear time as well.

### 6.2 Architecture details and hyperparameter settings

The hyperparameters used for our experiments are given in Table 1, Table 2, Table 3 and Table 4. An illustration of the sparsified linear mixing layers is given in Fig. 5.

**Table 1:** Hyperparameters used to train models on the rat LFP dataset for simulation.

| Hyperparameter | Value |
| --- | --- |
| In kernel size | 32 |
| Out kernel size | 32 |
| SC layers | 3 |
| SC kernel size | 32 |
| SC scales | 5 |
| Decay min | 1.0 |
| Decay max | 4.0 |
| Heads | 3 |
| Latent dim. per channel | 64 |
| Off-diag. size | 64 |
| Noise process | White noise |
| Schedule | Quadratic |
| Diffusion steps $T$ | 500 |
| B0 | 0.0001 |
| B1 | 0.06 |
| Time embedding dim. | 16 |
| Loss function | Mean-squared error (MSE) |
| Optimizer | AdamW |
| Learning rate | 0.0001 |
| Weight decay | 0.01 |
| Train batch size | 32 |
| Epochs | 500 |

**Table 2:** Hyperparameters used to train models on the 12 different AJILE12 datasets for our imputation experiment. The same hyperparameters were used for all 12 participants, with the exception of the number of training epochs, to account for the large differences in training data size between participants.

| Hyperparameter | Value |
| --- | --- |
| In kernel size | 1 |
| Out kernel size | 1 |
| SC layers | 3 |
| SC kernel size | 53 |
| SC scales | 4 |
| Decay min | 2.0 |
| Decay max | 2.0 |
| Heads | 3 |
| Latent dim. per channel | 16 |
| Off-diag. size | 4 |
| Noise process | OU with $\rho = 10$ |
| Schedule | Quadratic |
| Diffusion steps $T$ | 50 |
| B0 | 0.0001 |
| B1 | 0.5 |
| Time embedding dim. | 16 |
| Loss function | MSE |
| Optimizer | AdamW |
| Learning rate | 0.0005 |
| Weight decay | 0.01 |
| Train batch size | 32 |
| Epochs | 250/500/1000 |

**Table 3:** Hyperparameters used to train models on the macaque ECoG dataset for conditional generation of awake and anesthetized state.

| Hyperparameter | Value |
| --- | --- |
| In kernel size | 32 |
| Out kernel size | 32 |
| SC layers | 3 |
| SC kernel size | 101 |
| SC scales | 1 |
| Decay min | 2.0 |
| Decay max | 2.0 |
| Latent dim. per channel | 32 |
| Off-diag. size | 4 |
| Noise process | White noise |
| Schedule | Linear |
| Diffusion steps $T$ | 500 |
| B0 | 0.0001 |
| B1 | 0.03 |
| Time embedding dim. | 16 |
| Loss function | MSE |
| Optimizer | AdamW |
| Learning rate | 0.0004 |
| Weight decay | 0.01 |
| Train batch size | 32 |
| Epochs | 500 |

**Table 4:** Hyperparameters used to train models on the EEG BCI dataset.

| Hyperparameter | Value |
| --- | --- |
| In kernel size | 65 |
| Out kernel size | 65 |
| SC layers | 3 |
| SC kernel size | 65 |
| SC scales | 1 |
| Decay min | 2.0 |
| Decay max | 2.0 |
| Latent dim. per channel | 16 |
| Off-diag. size | 8 |
| Noise process | White noise |
| Schedule | Linear |
| Diffusion steps $T$ | 1000 |
| B0 | 0.0001 |
| B1 | 0.02 |
| Time embedding dim. | 16 |
| Loss function | MSE |
| Optimizer | AdamW |
| Learning rate | 0.0004 |
| Weight decay | 0.01 |
| Train batch size | 32 |
| Epochs | 1000 |

### 6.3 Additional results

Here we provide some additional results. In addition to PSDs, we can also compute cross-PSDs between different channels. For the LFP data recorded from rats, the median cross-PSDs match as well (Fig. 6).

**Figure 6:**
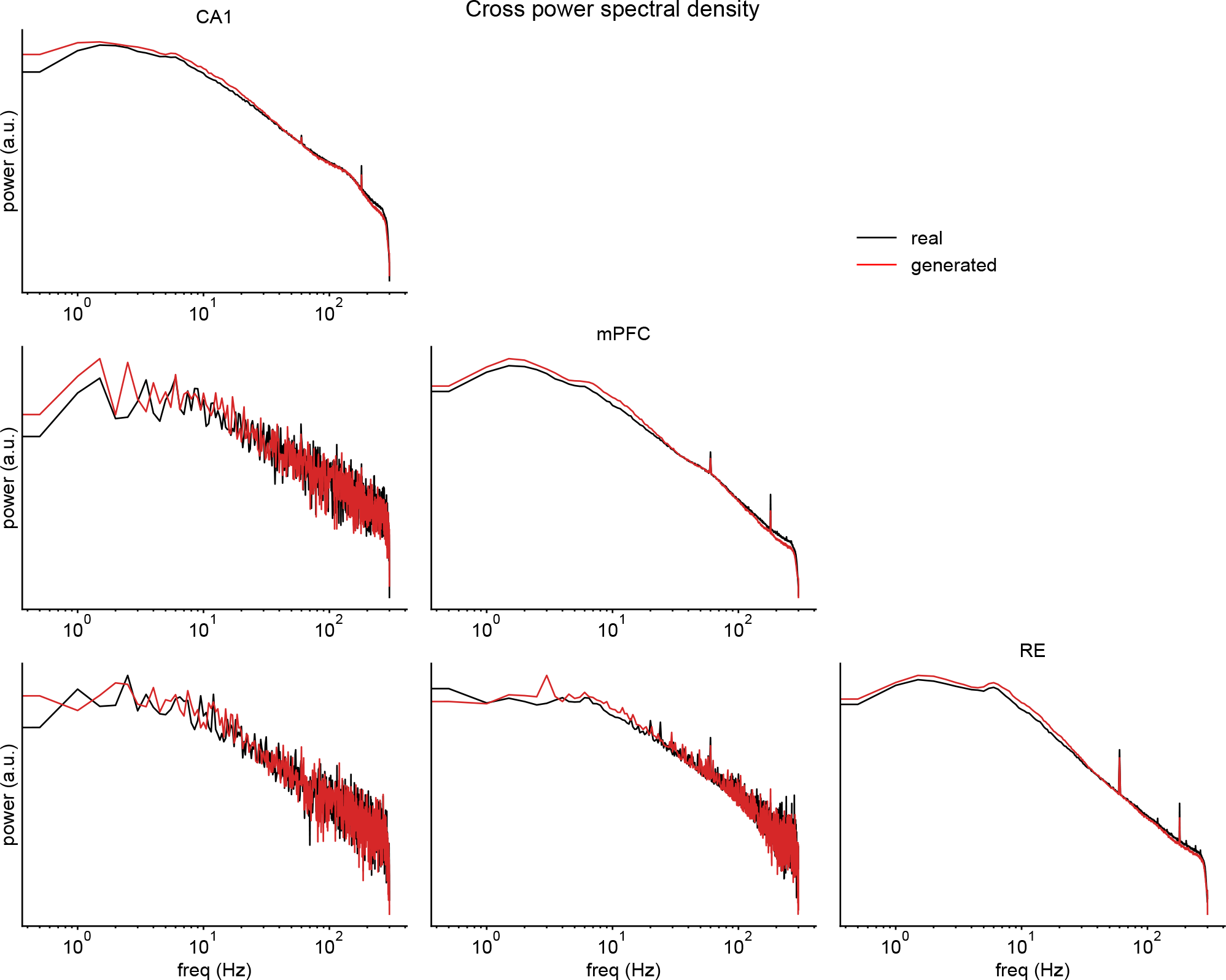
Median (cross-)PSDs for the rat LFP dataset for real and generated data.

Furthermore, for all datasets, the distribution of Euclidean distances between real and generated data is similar to the distribution of distances within. Crucially, the minimum distance between and within the real and generated time series is away from zero (Fig. 7). Therefore, the trained DDPMs do not overfit to the training data.

**Figure 7:**
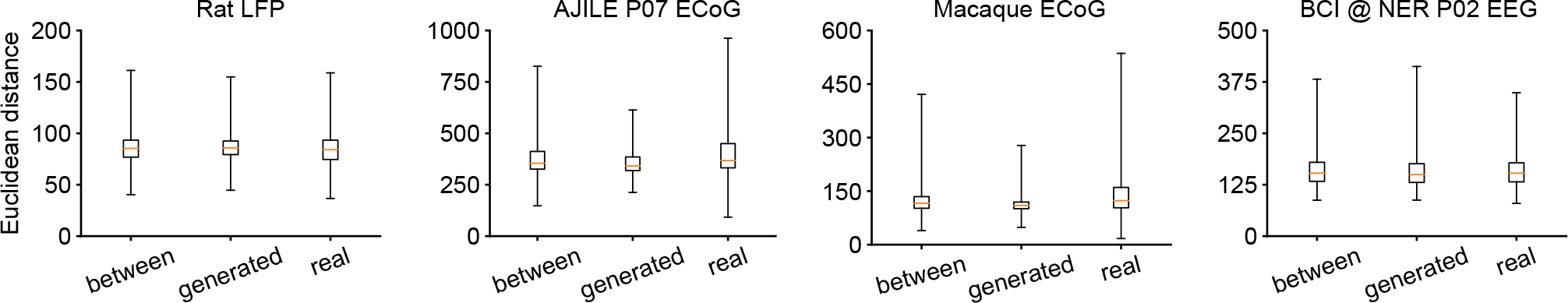
Distribution of distances between all real and generated time series, as well as within all real and generated time series. The box shows the median and 25% / 75% percentiles, and the whiskers show the minimum and maximum distance. The minimum distance between and within real and generated time series is away from zero.

We also provide all distributions of PSDs of AJILE12 participant P07 in the main text (Fig. 10). Additionally, all distributions of PSDs for two other participants, participant P01 with 94 channels (Fig. 11) and participant P12 with 126 channels (Fig. 12) are provided. In both cases, the median spectra as well as 10% and 90% percentiles match well. Recall that the neural decoding performance on data imputed by our model did not improve over the zero imputation baseline for participant P01. Nevertheless, the overall spectra of the real and imputed data match. This means that while the model was able to capture the overall distribution of time series, it failed to learn the class-conditional mapping between observed and missing channels.

**Figure 8:**
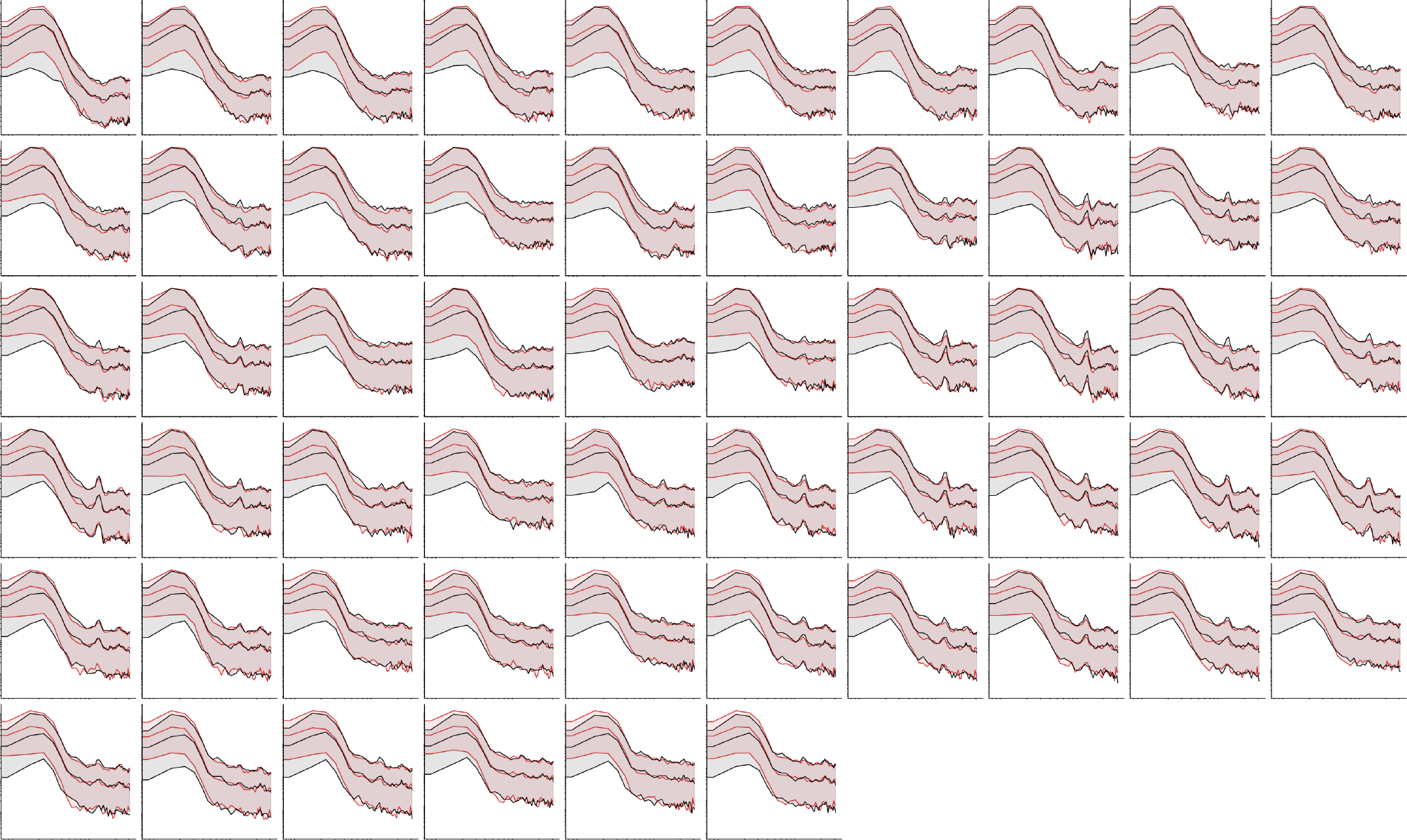
Full spectra of 56 channel EEG BCI data. Median power as well as 10% / 90% percentiles are shown.

**Figure 9:**
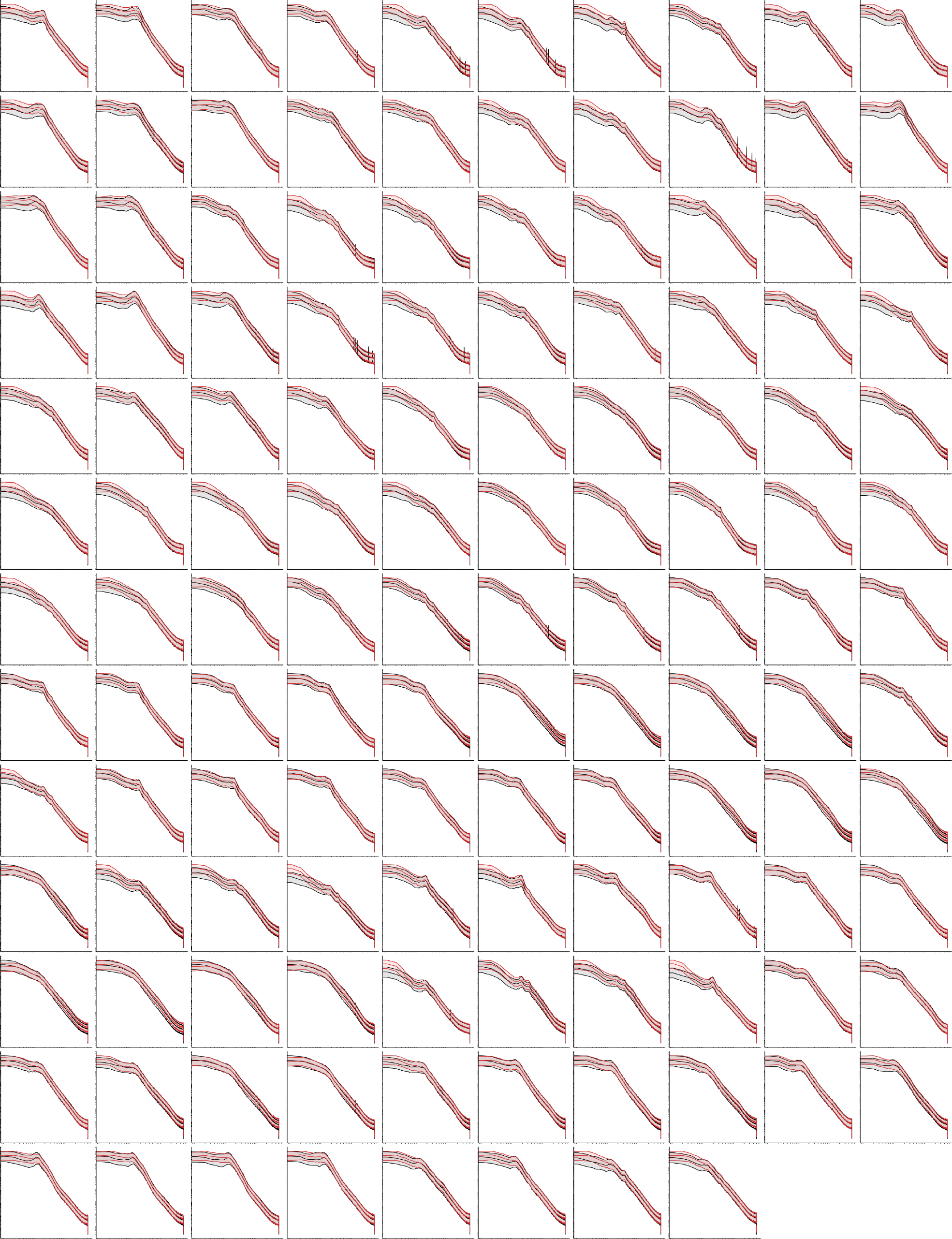
Full spectra of 128 channel macaque data in awake condition. Median power as well as 10% / 90% percentiles are shown.

**Figure 10:**
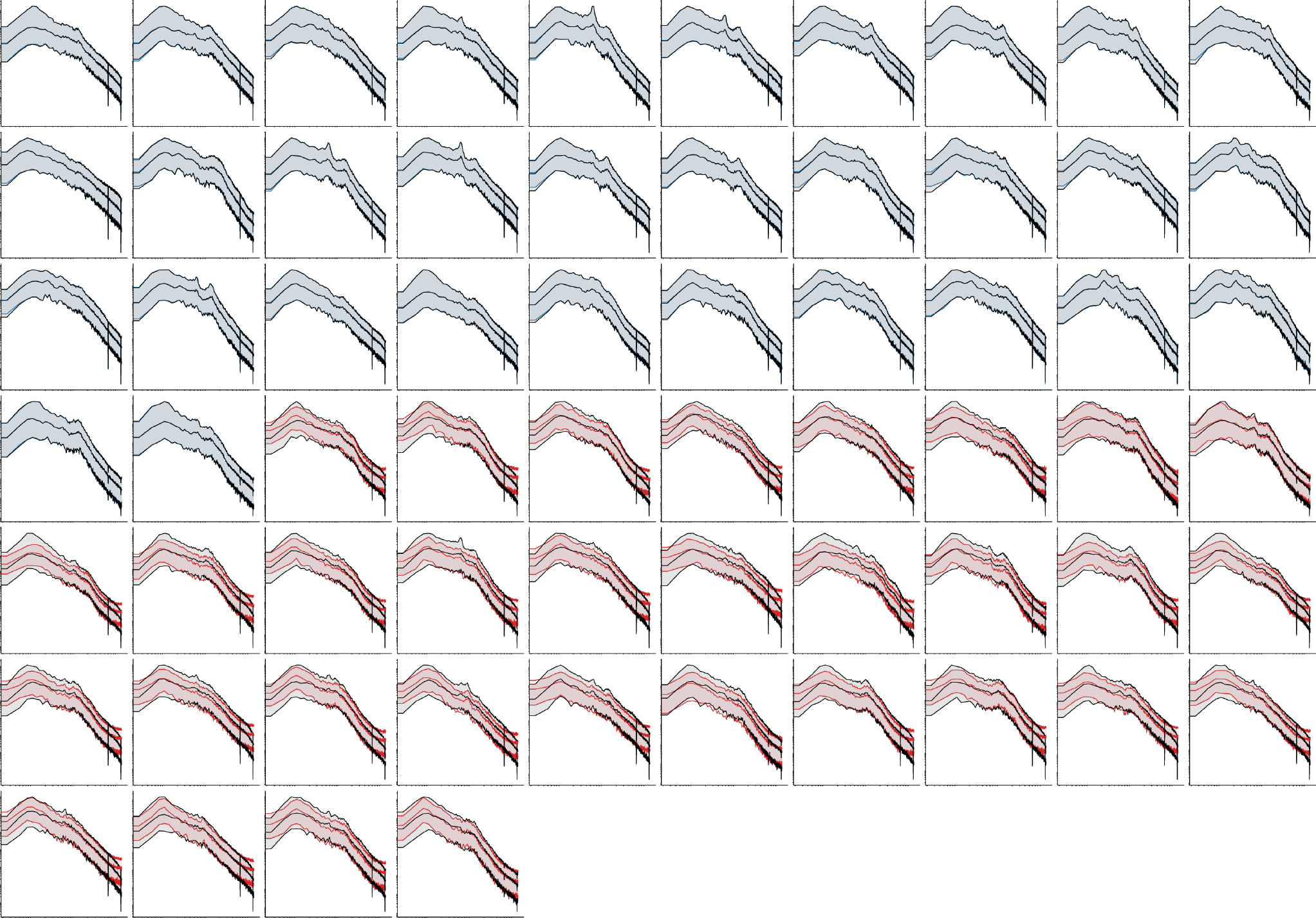
Full spectra of P07 from the AJILE12 dataset shown in the main text. The LFP traces consist of 64 channels. Here, we imputed the second half of the channels given the first half for all time series in the evaluation set. Median power as well as 10% / 90% percentiles are shown.

**Figure 11:**
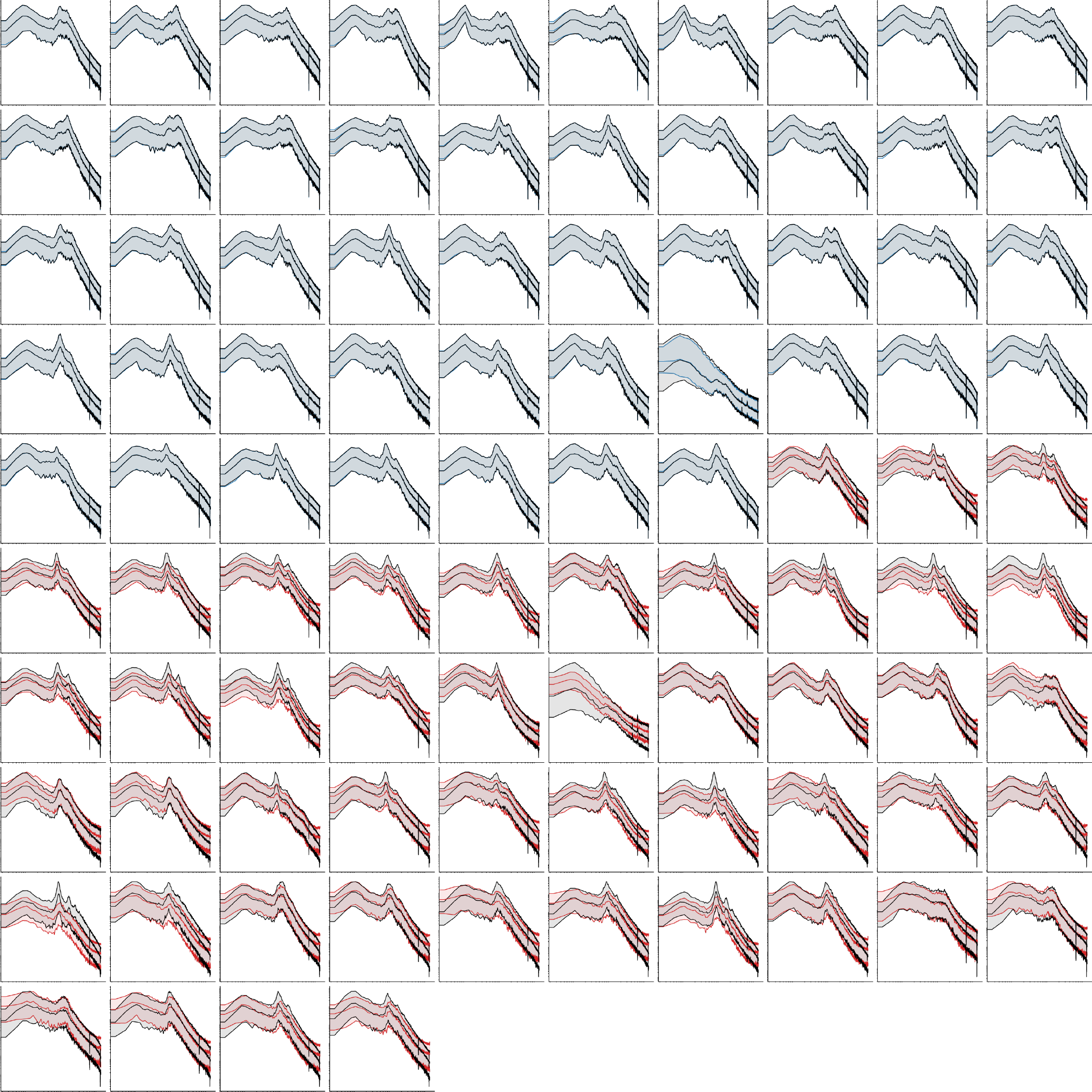
Full spectra of P01 from the AJILE12 dataset. The LFP traces consist of 94 channels. Here, we imputed the second half of the channels given the first half for all time series in the evaluation set. Median power as well as 10% / 90% percentiles are shown.

**Figure 12:**
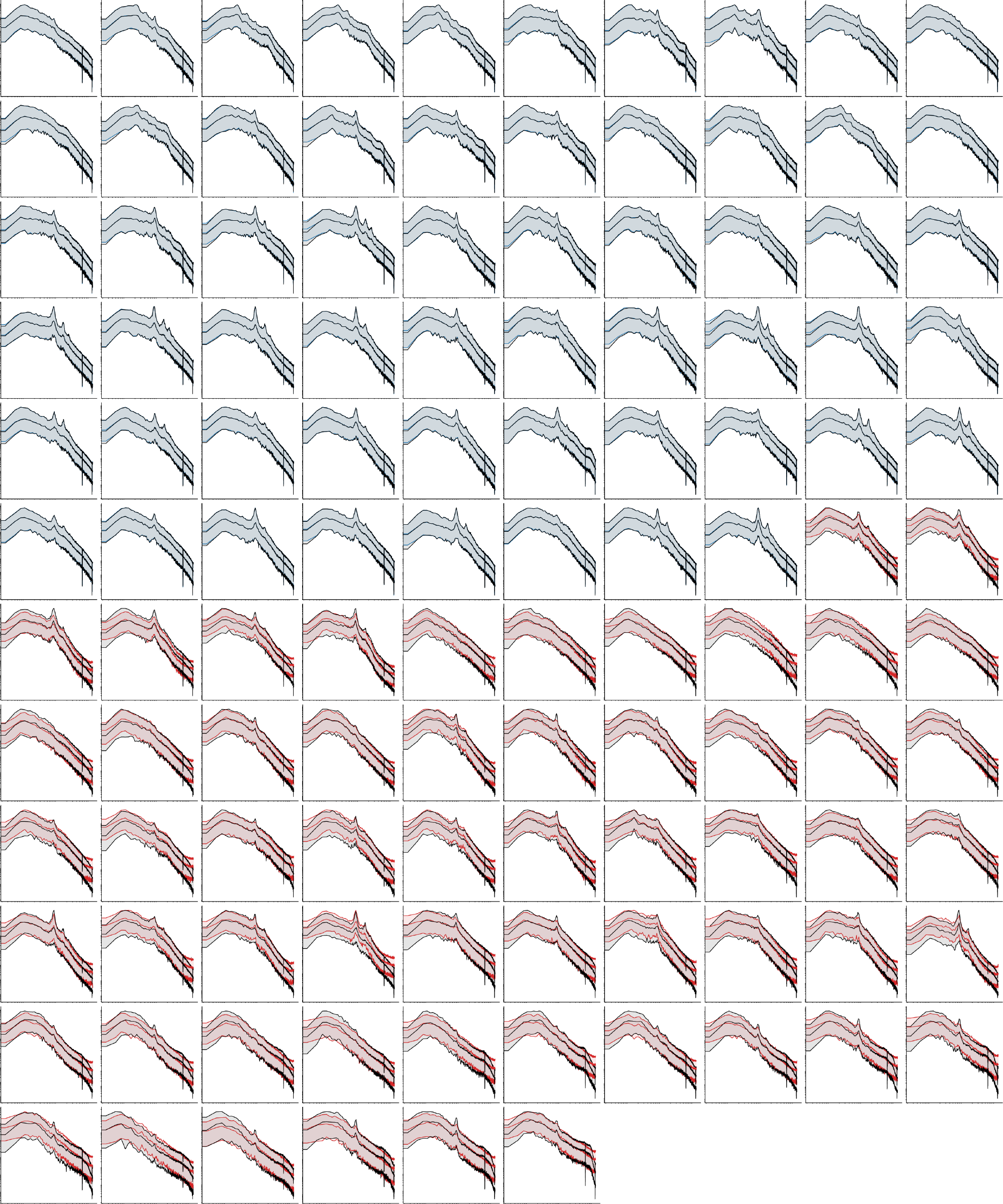
Full spectra of P12 from the AJILE12 dataset. The LFP traces consist of 126 channels. Here, we imputed the second half of the channels given the first half for all time series in the evaluation set. Median power as well as 10% / 90% percentiles are shown.

As an additional experiment with high channel count, we generated time series from the awake condition of the macaque ECoG data on all 128 recorded channels. The distribution of real and generated PSDs closely match across all channels, but our model sometimes overestimates the power in the low-frequency region (Fig. 9).

#### OU process versus white noise

Our use of the OU process provides a modification that can further improve the quality of generated samples in some cases. Here, we provide an analysis in terms of the spectral error for the AJILE12 dataset and the unconditional generation of the macaque ECoG data (awake only) on the full 128 channels. We retrain the models using the standard independent Gaussian process (white noise). All other training and network hyperparameters are kept fixed.

For both the (awake-only) macaque ECoG data and all AJILE12 participants, the spectral errors are significantly larger when using the white noise instead of the OU process (Fig. 13). This difference in performance is largely due to the low power, high frequency noise that is overestimated when white noise is used. However, for the rat LFP, the awake/anesthetized macaque ECoG data, and the BCI EEG data, there was no advantage to using the OU process over white noise.

**Figure 13:**
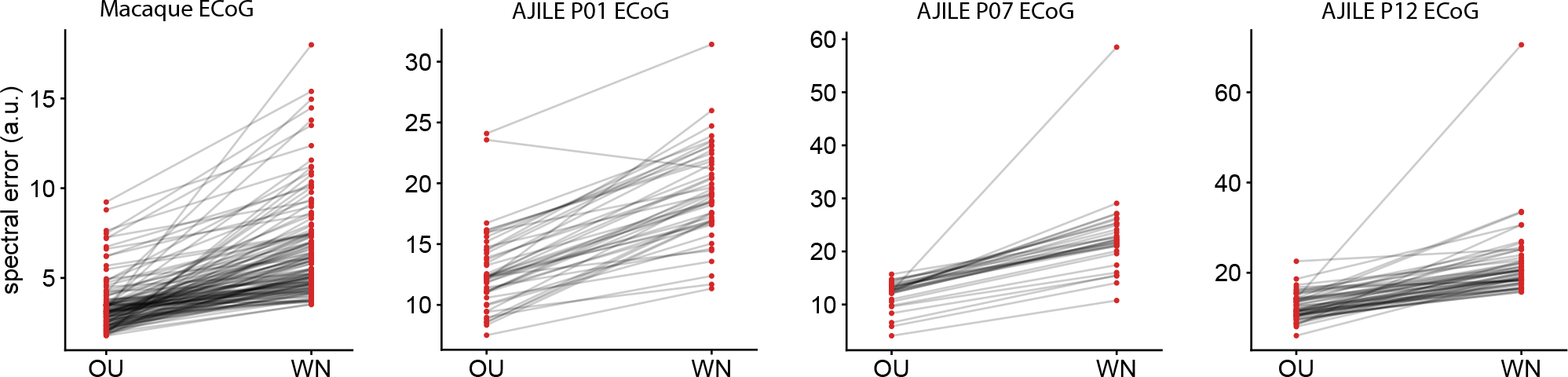
Comparison of OU versus white noise spectral errors of all generated / imputed channels for the macaque ECoG as well as AJILE12 participants P01, P07, and P12. All other training and network hyperparameters were fixed. The white noise process results in a higher spectral error.

## Footnotes

1 https://crcns.org/data-sets/hc/hc-24

2 https://dandiarchive.org/dandiset/000055/0.220127.0436

3 http://www.neurotycho.org/anesthesia-task

4 https://www.kaggle.com/competitions/inria-bci-challenge/

## References

Samira Abbasi, Selva Maran, and Dieter Jaeger. A general method to generate artificial spike train populations matching recorded neurons. Journal of Computational Neuroscience, 2020.

Juan Lopez Alcaraz and Nils Strodthoff. Diffusion-based time series imputation and forecasting with structured state space models. Transactions on Machine Learning Research, 2022.

Marin Biloš, Kashif Rasul, Anderson Schneider, Yuriy Nevmyvaka, and Stephan Günnemann. Modeling temporal data as continuous functions with process diffusion. arXiv preprint arXiv:2211.02590, 2022.

Kalok C Chan, G Andrew Karolyi, Francis A Longstaff, and Anthony B Sanders. An empirical comparison of alternative models of the short-term interest rate. The Journal of Finance, 1992.

Zijiao Chen, Jiaxin Qing, Tiange Xiang, Wan Lin Yue, and Juan Helen Zhou. Seeing beyond the brain: Conditional diffusion model with sparse masked modeling for vision decoding. Proceedings of the IEEE/CVF Conference on Computer Vision and Pattern Recognition, 2023.

Prafulla Dhariwal and Alexander Nichol. Diffusion models beat GANs on image synthesis. Advances in Neural Information Processing Systems, 2021.

Thomas Donoghue, Matar Haller, Erik J Peterson, Paroma Varma, Priyadarshini Sebastian, Richard Gao, Torben Noto, Antonio H Lara, Joni D Wallis, Robert T Knight, Avgusta Shestyuk, and Bradley Voytek. Parameterizing neural power spectra into periodic and aperiodic components. Nature Neuroscience, 2020.

Lawrence Ashley Farwell and Emanuel Donchin. Talking off the top of your head: Toward a mental prosthesis utilizing event-related brain potentials. Electroencephalography and clinical Neurophysiology, 1988.

Vincent Fortuin, Dmitry Baranchuk, Gunnar R ätsch, and Stephan Mandt. GP-VAE: Deep probabilistic time series imputation. International Conference on Artificial Intelligence and Statistics, 2020.

Cecilia Gallego-Carracedo, Matthew G Perich, Raeed H Chowdhury, Lee E Miller, and Juan Álvaro Gallego. Local field potentials reflect cortical population dynamics in a region-specific and frequency-dependent manner. eLife, 2022.

Richard Gao, Erik J Peterson, and Bradley Voytek. Inferring synaptic excitation/inhibition balance from field potentials. NeuroImage, 2017.

Alessandro T Gifford, Kshitij Dwivedi, Gemma Roig, and Radoslaw M Cichy. A large and rich EEG dataset for modeling human visual object recognition. NeuroImage, 2022.

Albert Gu, Karan Goel, and Christopher Ré. Efficiently modeling long sequences with structured state spaces. International Conference on Learning Representations, 2022.

Diego A Gutnisky and Krešimir Josić. Generation of spatiotemporally correlated spike trains and local field potentials using a multivariate autoregressive process. Journal of Neurophysiology, 2010.

Charles R Harris, K Jarrod Millman, Stéfan J Van Der Walt, Ralf Gommers, Pauli Virtanen, David Cour-napeau, Eric Wieser, Julian Taylor, Sebastian Berg, Nathaniel J Smith, et al. Array programming with NumPy. Nature, 2020.

Biyu J He, John M Zempel, Abraham Z Snyder, and Marcus E Raichle. The temporal structures and functional significance of scale-free brain activity. Neuron, 2010.

Dan Hendrycks and Kevin Gimpel. Gaussian error linear units. arXiv preprint arXiv:1606.08415, 2016.

Jonathan Ho, Ajay Jain, and Pieter Abbeel. Denoising diffusion probabilistic models. Advances in Neural Information Processing Systems, 2020.

Sergey Ioffe and Christian Szegedy. Batch normalization: Accelerating deep network training by reducing internal covariate shift. International Conference on Machine Learning, pages 448–456, 2015.

Tero Karras, Miika Aittala, Timo Aila, and Samuli Laine. Elucidating the design space of diffusion-based generative models. Advances in Neural Information Processing Systems, 2022.

Zhifeng Kong, Wei Ping, Jiaji Huang, Kexin Zhao, and Bryan Catanzaro. Diffwave: A versatile diffusion model for audio synthesis. International Conference on Learning Representations, 2021.

Michael Krumin and Shy Shoham. Generation of spike trains with controlled auto-and cross-correlation functions. Neural Computation, 2009.

Yann LeCun, Yoshua Bengio, et al. Convolutional networks for images, speech, and time series. The handbook of brain theory and neural networks, 1995.

Yuhong Li, Tianle Cai, Yi Zhang, Deming Chen, and Debadeepta Dey. What makes convolutional models great on long sequence modeling? International Conference on Learning Representations, 2023.

Lequan Lin, Zhengkun Li, Ruikun Li, Xuliang Li, and Junbin Gao. Diffusion models for time series applications: A survey. arXiv preprint arXiv:2305.00624, 2023.

Sikun Lin, Thomas Sprague, and Ambuj K Singh. Mind reader: Reconstructing complex images from brain activities. Advances in Neural Information Processing Systems, 2022.

Ilya Loshchilov and Frank Hutter. Decoupled weight decay regularization. International Conference on Learning Representations, 2019.

Cheng Lu, Yuhao Zhou, Fan Bao, Jianfei Chen, Chongxuan Li, and Jun Zhu. DPM-Solver: A fast ODE solver for diffusion probabilistic model sampling in around 10 steps. Advances in Neural Information Processing Systems, 2022.

Andreas Lugmayr, Martin Danelljan, Andres Romero, Fisher Yu, Radu Timofte, and Luc Van Gool. Repaint: Inpainting using denoising diffusion probabilistic models. Proceedings of the IEEE/CVF Conference on Computer Vision and Pattern Recognition, 2022.

Jakob H Macke, Philipp Berens, Alexander S Ecker, Andreas S Tolias, and Matthias Bethge. Generating spike trains with specified correlation coefficients. Neural Computation, 2009.

Perrin Margaux, Maby Emmanuel, Daligault Sébastien, Bertrand Olivier, and Mattout Jérémie. Objective and subjective evaluation of online error correction during P300-based spelling. Advances in Human-Computer Interaction, 2012.

Manuel Molano-Mazon, Arno Onken, Eugenio Piasini, and Stefano Panzeri. Synthesizing realistic neural population activity patterns using generative adversarial networks. International Conference on Learning Representations, 2018.

Markus Ojala and Gemma C. Garriga. Permutation tests for studying classifier performance. Journal of Machine Learning Research, 2010.

Chethan Pandarinath, Daniel J O’Shea, Jasmine Collins, Rafal Jozefowicz, Sergey D Stavisky, Jonathan C Kao, Eric M Trautmann, Matthew T Kaufman, Stephen I Ryu, Leigh R Hochberg, et al. Inferring singletrial neural population dynamics using sequential auto-encoders. Nature Methods, 2018.

Adam Paszke, Sam Gross, Francisco Massa, Adam Lerer, James Bradbury, Gregory Chanan, Trevor Killeen, Zeming Lin, Natalia Gimelshein, Luca Antiga, et al. Pytorch: An imperative style, high-performance deep learning library. Advances in neural information processing systems, 2019.

Steven M Peterson, Zoe Steine-Hanson, Nathan Davis, Rajesh PN Rao, and Bingni W Brunton. Generalized neural decoders for transfer learning across participants and recording modalities. Journal of Neural Engineering, 2021.

Steven M Peterson, Satpreet H Singh, Benjamin Dichter, Michael Scheid, Rajesh PN Rao, and Bingni W Brunton. AJILE12: Long-term naturalistic human intracranial neural recordings and pose. Scientific data, 2022.

Carlos R Ponce, Will Xiao, Peter F Schade, Till S Hartmann, Gabriel Kreiman, and Margaret S Livingstone. Evolving images for visual neurons using a deep generative network reveals coding principles and neuronal preferences. Cell, 2019.

W S Pritchard. The brain in fractal time: 1/f-like power spectrum scaling of the human electroencephalogram. The International journal of neuroscience, 1992.

Poornima Ramesh, Mohamad Atayi, and Jakob H Macke. Adversarial training of neural encoding models on population spike trains. Real Neurons & Hidden Units: Future directions at the intersection of neuroscience and artificial intelligence @ NeurIPS 2019, 2019.

Kashif Rasul, Calvin Seward, Ingmar Schuster, and Roland Vollgraf. Autoregressive denoising diffusion models for multivariate probabilistic time series forecasting. International Conference on Machine Learning, 2021.

Robin Rombach, Andreas Blattmann, Dominik Lorenz, Patrick Esser, and Björn Ommer. High-resolution image synthesis with latent diffusion models. Proceedings of the IEEE/CVF Conference on Computer Vision and Pattern Recognition, 2022.

Benjamin Sanchez-Lengeling and Alán Aspuru-Guzik. Inverse molecular design using machine learning: Generative models for matter engineering. Science, 2018.

Thomas Schreiber and Andreas Schmitz. Surrogate time series. Physica D: Nonlinear Phenomena, 2000.

Ikaro Silva, George Moody, Daniel J Scott, Leo A Celi, and Roger G Mark. Predicting in-hospital mortality of icu patients: The physionet/computing in cardiology challenge 2012. Computing in Cardiology, 2012.

Jiaming Song, Chenlin Meng, and Stefano Ermon. Denoising diffusion implicit models. International Conference on Learning Representations, 2021a.

Yang Song, Jascha Sohl-Dickstein, Diederik P Kingma, Abhishek Kumar, Stefano Ermon, and Ben Poole. Score-based generative modeling through stochastic differential equations. International Conference on Learning Representations, 2021b.

He Sun and Katherine L Bouman. Deep probabilistic imaging: Uncertainty quantification and multi-modal solution characterization for computational imaging. Proceedings of the AAAI Conference on Artificial Intelligence, 2021.

Yu Takagi and Shinji Nishimoto. High-resolution image reconstruction with latent diffusion models from human brain activity. bioRxiv, 2022.

Sabera Talukder, Jennifer J Sun, Matthew Leonard, Bingni W Brunton, and Yisong Yue. Deep neural imputation: A framework for recovering incomplete brain recordings. NeurIPS 2022 Workshop on Learning from Time Series for Health, 2022.

Yusuke Tashiro, Jiaming Song, Yang Song, and Stefano Ermon. CSDI: Conditional score-based diffusion models for probabilistic time series imputation. Advances in Neural Information Processing Systems, 2021.

Gerrit van den Burg and Chris Williams. On memorization in probabilistic deep generative models. Advances in Neural Information Processing Systems, 2021.

Carmen Varela and Matthew A Wilson. Simultaneous extracellular recordings from midline thalamic nuclei, medial prefrontal cortex and CA1 from rats cycling through bouts of sleep and wakefulness. CRCNS.org, 2019.

Carmen Varela and Matthew A Wilson. mPFC spindle cycles organize sparse thalamic activation and recently active CA1 cells during non-REM sleep. eLife, 2020.

Victor Venema, Felix Ament, and Clemens Simmer. A stochastic iterative amplitude adjusted Fourier transform algorithm with improved accuracy. Nonlinear Processes in Geophysics, 2006.

Pauli Virtanen, Ralf Gommers, Travis E Oliphant, Matt Haberland, Tyler Reddy, David Cournapeau, Evgeni Burovski, Pearu Peterson, Warren Weckesser, Jonathan Bright, et al. SciPy 1.0: Fundamental algorithms for scientific computing in python. Nature Methods, 2020.

Saurabh Vyas, Matthew D Golub, David Sussillo, and Krishna V Shenoy. Computation through neural population dynamics. Annual Review of Neuroscience, 2020.

Jason Walonoski, Mark Kramer, Joseph Nichols, Andre Quina, Chris Moesel, Dylan Hall, Carlton Duffett, Kudakwashe Dube, Thomas Gallagher, and Scott McLachlan. Synthea: An approach, method, and software mechanism for generating synthetic patients and the synthetic electronic health care record. Journal of the American Medical Informatics Association, 2018.

Zhiguang Wang, Weizhong Yan, and Tim Oates. Time series classification from scratch with deep neural networks: A strong baseline. 2017 International joint conference on neural networks (IJCNN), 2017.

Toru Yanagawa, Zenas C Chao, Naomi Hasegawa, and Naotaka Fujii. Large-scale information flow in conscious and unconscious states: An ECoG study in monkeys. PloS one, 2013.

Jinsung Yoon, Daniel Jarrett, and Mihaela Van der Schaar. Time-series generative adversarial networks. Advances in Neural Information Processing Systems, 2019.

